# Erosion of the Epigenetic Landscape and Loss of Cellular Identity as a Cause of Aging in Mammals

**DOI:** 10.1101/808642

**Authors:** Jae-Hyun Yang, Patrick T. Griffin, Daniel L. Vera, John K. Apostolides, Motoshi Hayano, Margarita V. Meer, Elias L. Salfati, Qiao Su, Elizabeth M. Munding, Marco Blanchette, Mital Bhakta, Zhixun Dou, Caiyue Xu, Jeffrey W. Pippin, Michael L. Creswell, Brendan L. O’Connell, Richard E. Green, Benjamin A. Garcia, Shelley L. Berger, Philipp Oberdoerffer, Stuart J. Shankland, Vadim N. Gladyshev, Luis A. Rajman, Andreas R. Pfenning, David A. Sinclair

## Abstract

All living things experience entropy, manifested as a loss of inherited genetic and epigenetic information over time. As budding yeast cells age, epigenetic changes result in a loss of cell identity and sterility, both hallmarks of yeast aging. In mammals, epigenetic information is also lost over time, but what causes it to be lost and whether it is a cause or a consequence of aging is not known. Here we show that the transient induction of genomic instability, in the form of a low number of non-mutagenic DNA breaks, accelerates many of the chromatin and tissue changes seen during aging, including the erosion of the epigenetic landscape, a loss of cellular identity, advancement of the DNA methylation clock and cellular senescence. These data support a model in which a loss of epigenetic information is a cause of aging in mammals.

**One Sentence Summary:** The act of repairing DNA breaks induces chromatin reorganization and a loss of cell identity that may contribute to mammalian aging

## INTRODUCTION

The preservation of information is essential for any ordered system to endure, including life. Living things use two main information storage systems: genetic and epigenetic. In the face of considerable chaos at the nanoscale, life endures by utilizing energy from its environment to preserve information. Over time, however, information laid down during development is ultimately lost. In 1948, Claude Shannon’s Mathematical Theory of Communication proposed methods to accurately transfer information over time and space, forming the basis of today’s global communication networks but may also be relevant to aging (Sinclair and LaPlante, 2019).

In the 1950’s, Medawar and Szilard independently proposed that aging is due to a loss of information caused by mutations (Medawar, 1952; Szilard, 1959). Evidence for hypothesis included the fact that mutations accumulate (Martincorena et al., 2018; Martincorena et al., 2015; Zhang et al., 2019) and increased genome instability seems to accelerate aging in mice and humans (Huang et al., 2006; Li et al., 2007; Niedernhofer et al., 2006; Tan et al., 2005). In the past decade, however, conflicting data has prompted a reevaluation of this hypothesis. For example, certain strains of mice engineered to accumulate mutations at a faster rate do not experience accelerated aging (progeria) (Narayanan et al., 1997), while others with a reduced DNA repair efficiency experience progeria without an increase in mutation frequency (Dolle et al., 2006). And in aged mice, defects in mitochondrial function can be reversed within a few days, indicating that mutations in mitochondrial DNA are unlikely to cause aging until late in life (Gomes et al., 2013). But one of the most difficult observations to reconcile with the hypothesis is somatic cell nuclear transfer, which can generate individuals of a variety of mammalian species that are healthy and have normal lifespans (Burgstaller and Brem, 2017). Thus, while DNA damage and mutation accumulation are both associated with aging, these finding point to aging being the result of another, closely-related process.

Around the same time as Medawar and Szilard, Waddington proposed the concept of the epigenetic landscape, a metaphor for the process by which cells with identical genomes give rise to different cell types (Waddington, 1957). Today, we understand that cellular differentiation and specialization is largely governed by transcriptional networks and epigenetic modifications that allow for information to be passed on during replication and stored for the life of the cell and its progeny. In the 1990s, studies of budding yeast provided the first evidence that an erosion of and “smoothening” of the epigenetic landscape may underlie aspects of aging (Imai and Kitano, 1998; Kennedy et al., 1995; Kennedy et al., 1997; Sinclair and Guarente, 1997; Sinclair et al., 1997).

Young yeast cells are fertile because they only express genes from one gender, either **a** or α mating type information. The repositories of **a** and α information at the silent *HM* loci are kept that way by the silent information regulator complex, the catalytic component of which is Sir2, an NAD^+^-dependent histone deacetylase (Imai et al., 2000). As yeast cells age, double-stranded DNA breaks (DSBs) initiate a DNA damage signal that recruits Sir2 and other epigenetic factors, such as Hst1, Rpd3, Gcn5, and Esa1, to sites of DNA repair (Martin et al., 1999; McAinsh et al., 1999; Mills et al., 1999; Tamburini and Tyler, 2005). After repair, these factors return to their original sites, but as the number of DSBs and genome instability increases, Sir2 is increasingly sequestered away from *HM* loci, causing cells lose their identity and become sterile, a defining hallmark of yeast aging (Smeal et al., 1996). These findings led us to propose a model in which the relocalization of chromatin modifiers (RCM) introduces non-random changes to the epigenome that cause aging (Oberdoerffer et al., 2008).

Like yeast and other more complex eukaryotes (Benayoun et al., 2015), mammals also undergo a characteristic set of epigenetic changes as they age (Kane and Sinclair, 2019), including a decrease in the abundance of histones H3 and H4 and the heterochromatin marks H3K9me3 and H3K27me3, along with an overall decrease in DNA methylation (DNAme) at CpG sites (Cruickshanks et al., 2013; O’Sullivan et al., 2010; Scaffidi and Misteli, 2006; Shumaker et al., 2006; Singhal et al., 1987; Zhang et al., 2015). Site-specific DNA methylation changes also occur over time, forming the basis of “epigenetic clocks” (Hannum et al., 2013; Horvath, 2013; Meer et al., 2018; Petkovich et al., 2017; Stubbs et al., 2017; Wang et al., 2017). With age, the heterogeneity of both transcriptional patterns chromatin modifications in various cell types, including cardiomyocytes, CD4 T-cells, pancreatic cells and skin, also increases (Bahar et al., 2006; Martinez-Jimenez et al., 2017; Salzer et al., 2018), consistent with an increase in “informational or epigenetic entropy” (Field et al., 2018; Hannum et al., 2013; Jenkinson et al., 2017).

The mammalian RCM response is thought to drive aging in a similar way to yeast (Oberdoerffer et al., 2008). Three mammalian Sir2 homologs (sirtuins), namely SIRT1, SIRT6, and SIRT7, also associate with rDNA to promote the formation of heterochromatin. SIRT1 and SIRT6 are also bound to loci involved in regulating inflammation, growth, energy metabolism, and in suppressing LINE-1 elements (Simon et al., 2019). SIRT7, meanwhile, promotes genome stability at the rDNA locus, forestalling cellular senescence (Paredes et al., 2018). In response to a DNA damage signal, one of which is known to be ATM (Dobbin et al., 2013; Oberdoerffer et al., 2008), sirtuins relocalize to DSB where they promote repair, ostensibly by deacetylating histones, other chromatin modifiers (e.g. HDAC1) and DNA repair proteins (e.g. PARP1, SNF2H and NBS1) (Dobbin et al., 2013; Mao et al., 2011; Oberdoerffer et al., 2008; Toiber et al., 2013). The ATM-dependent relocalization of SIRT1 to DSBs, for example, is associated with increased H1K26 acetylation, the ectopic expression of SIRT1-repressed genes, and the ectopic transcription of satellite and LINE-1 elements (Oberdoerffer et al., 2008). Consistent with the RCM hypothesis, SIRT6 activity and DSB repair efficiency, but not other types of DNA repair, correlate with the maximum lifespan of diverse mammalian species (Tian et al., 2019).

Similar to yeast aging, an increasing number of studies indicate that a dysregulation of developmental genes that control cell identity is associated with mammalian aging. In mice, for example, hematopoietic stem cells switch from canonical to non-canonical Wnt signaling (Florian et al., 2013), old dermal fibroblasts gain adipogenic traits (Salzer et al., 2018) and muscle satellite cells activate Wnt signaling, converting them from a myogenic to a fibrogenic lineage (Brack et al., 2007). Satellite cells also activate *Hoxa9*, which activates Wnt, TGFβ and JAK/STAT signaling in response to BaCl2-induced muscle injury (Schworer et al., 2016). In aging human brain tissue, too, there is a distinct upregulation of developmental genes (Donertas et al., 2018).

These and other observations indicate that perturbations that reorganize the epigenome, such as DSBs, induce transcriptional and epigenetic noise during aging (Bahar et al., 2006; Wiley et al., 2017). But whether or not these changes are a cause of mammalian aging or merely a consequence is the subject of considerable speculation. To test this, we generated a system called *ICE* (for inducible <u>c</u>hanges to the <u>e</u>pigenome), which allows for precise, non-mutagenic DSBs to be created in cultured cells and in mice. Using this system, we provide evidence that DSBs induce epigenomic noise that erodes the epigenetic landscape in predictable ways, leading to a loss of cellular identity, advancement of the DNAme clock, and an age-related decline in function, which together support the hypothesis that a loss of epigenetic information over time is a conserved cause of aging in eukaryotes.

## RESULTS

### A System to Induce Epigenetic Aging

We sought to test the possibility a smoothening of the epigenetic landscape is a driver of age-related molecular and physiological change (Jenkinson et al., 2017; Kennedy et al., 1995; Oberdoerffer et al., 2008). In this model, DSBs repeatedly recruit chromatin modifiers to the break site to mediate repair but, over time, incomplete chromatin restoration causes epigenetic information to be lost, leading to alterations in gene expression and susceptibility to more DSBs (**Figure 1A**).

**Figure 1.**
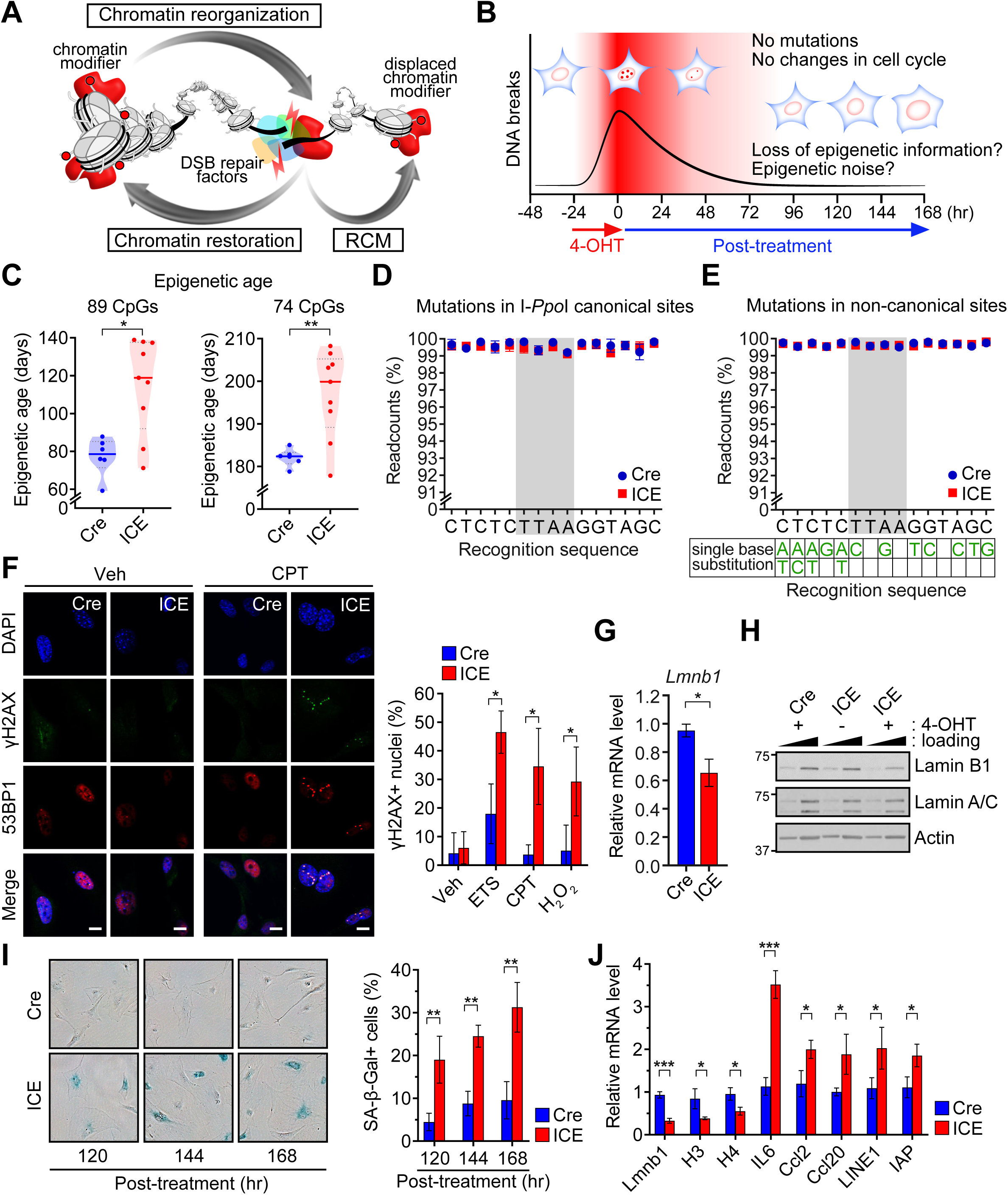
A Cell-based System to Study Mechanisms of Epigenetic Aging. (A) The RCM hypothesis states that a loss of epigenetic information due to chromatin disruptors such as DSBs accelerate aging. (B) Time course of DSB induction by 4-Hydroxytamoxifen (4-OHT) and changes that occur after ICE cells have completed mutation-free DNA repair. (C) Epigenetic age of ICE cells post-treatment. All DNA methylation sites (left) and DNA methylation sites post-batch effect correction (right). Mann-Whitney U test. (D and E) Percent non-mutated I-*Ppo*I canonical and non-canonical recognition sequences in post-treated ICE cells assessed by deep sequencing (>50x). (F) Immunostaining of DNA damage markers γH2AX and 53BP1 in post-treated ICE cells with and without exposure to the DNA damaging agents (ETS, etoposide; CPT, camptothecin; H_2_O_2_, hydrogen peroxide. Scale bar, 10 µm. Two-tailed Student’s *t* test. (G and H) Lamin B1 mRNA and protein levels in post-treated ICE cells. Lamin A/C and actin are loading controls known not to change during senescence. (I) Images and quantification of SA-β-Gal staining of post-treated ICE and Cre cells. Two-tailed Student’s *t* test. (J) mRNA levels of genes known to change during senescence at 144hr post-treatment. Two-tailed Student’s *t* test. Data are mean (n≥3) ± SD. *p < 0.05; **p < 0.01; ***p< 0.001.

Due to the complexity of mammals, however, studies of this process, including our own (Oberdoerffer et al., 2008), have been largely correlative. To test cause and effect, we created the ICE mouse strain with a tamoxifen-inducible Cre and a I-*Ppo*I homing endonuclease that recognizes ∼20 sites in the genome, although not all are accessible (Berkovich et al., 2007; Kim et al., 2016). This allowed us to cut the genome in specific places and induce epigenomic noise in a way that mimicked naturally occurring DSBs (Berkovich et al., 2007; van Sluis and McStay, 2015). In the accompanying paper, we show that the ICE system allows for precise temporal and spatial control over DSBs, that DSBs are induced predominately in non-coding regions at frequencies only a few-fold above background, and cutting does not cause cell cycle pausing, senescence, or mutations (Hayano et al, co-submitted manuscript). Triggering of the ICE system for a few weeks in young animals accelerates the pace of aging at the molecular, physiological and histological levels, including a 50% acceleration of the DNA methylation clock.

To understand the molecular processes that drive epigenetic aging, we developed an *in vitro* system consisting of murine embryonic fibroblasts (MEFs) from ICE mice (Cre^−/+^/I-*Ppo*I^−/+^) and controls (Cre^−/+^/I-*Ppo*I^−/−^). We used MEFs rather than adult cells because of their lower epigenomic entropy and cells from embryos of the same litter to ensure they were the same age and had similar transcription profiles (**Figure S1A**). Growth rate and cell cycle kinetics were also similar between all isolates prior to induction. I-*Ppo*I was then induced for 24 hours by incubating cells in the presence of 4-Hydroxytamoxifen (4-OHT), leading to ∼4-fold increase in γH2AX foci over background, with no evidence of cell cycle changes, senescence, telomere shortening, or genome instability during or after treatment (**Figure 1B**) (also see Hayano et al, co-submitted manuscript).

The ICE mice experienced epigenetic aging faster than their age-matched counterparts. To test if this effect is cell-intrinsic, three independent ICE cell lines were treated as above and their DNA was analyzed by reduced representation bisulfite sequencing (RRBS). Using a weighted average of 89 age-associated methylation sites, or a refined set of 74 sites (Petkovich et al., 2017), the DNA methylation age of the cells was on average ∼1.5-fold greater than the Cre control cells, closely paralleling the ICE mice (**Figure S1B and Figure 1C;** p=0.01, p=0.008, respectively).

Based on extensive whole-genome sequencing, there was no difference in the number of mutations in Cre and ICE cells after cutting and recovery for 96 hours, and no difference in mutation frequency at canonical, non-canonical, or 100,000 random sites (**Figure 1D**, **1E**, **S1C and S1D**). The same result was seen in skeletal muscle from ICE mice (Hayano et al., co-submitted manuscript).

Next, we tested if the epigenetically-aged ICE cells displayed characteristics of cells from old mice. One of the most robust and reproducible effects of aging is an increased sensitivity to DNA damaging agents including camptothecin, etoposide, and hydrogen peroxide (Li et al., 2016; Mapuskar et al., 2017; Miyoshi et al., 2006). After recovery from cutting ICE cells were significantly more susceptible than Cre controls to DNA damage caused by the above agents, based on increased numbers of γH2AX and 53BP1 foci (**Figure 1F**, **S1E and S1F**).

Another hallmark of aging is a decrease in Lamin B1 that promotes cellular senescence, as indicated by SA-β-Gal activity and markers such as IL6, Ccl2, Ccl20, LINE-1 and IAP (Freund et al., 2012; Shah et al., 2013). At later time points after I-*Ppo*I induction (96-168 hrs), ICE cell cultures had lower levels of Lamin B1 (**Figure 1G**, **1H and S1G**) and increased proportions of cells that were ostensibly senescent (**Figure 1I and 1J**).

### DNA Breaks Erode the Epigenetic Landscape

Having established that non-mutagenic DSBs accelerate the DNA methylation clock in cell culture and in mice, we set out to explore the underlying mechanisms. To assess global epigenetic changes, histones were purified from Cre and ICE cells 96 hours post-induction and a total of 46 different post-translational histone modifications were quantified using mass spectrometry (MSMS). There were three significant changes: a decrease in the acetylation of histones H3 at lysines 27 and 56 (H3K27ac and H3K56ac) and an increase in acetylation of H3 at lysine 122 (H3K122ac) (**Figure 2A**, **2B**). Interestingly, H3K27ac and H3K56ac are decreased in many human immune cell types at the single cell level (Cheung et al., 2018; Dang et al., 2009) and reduced levels of H3K122ac extend the lifespan of yeast (Sen et al., 2015).

**Figure 2.**
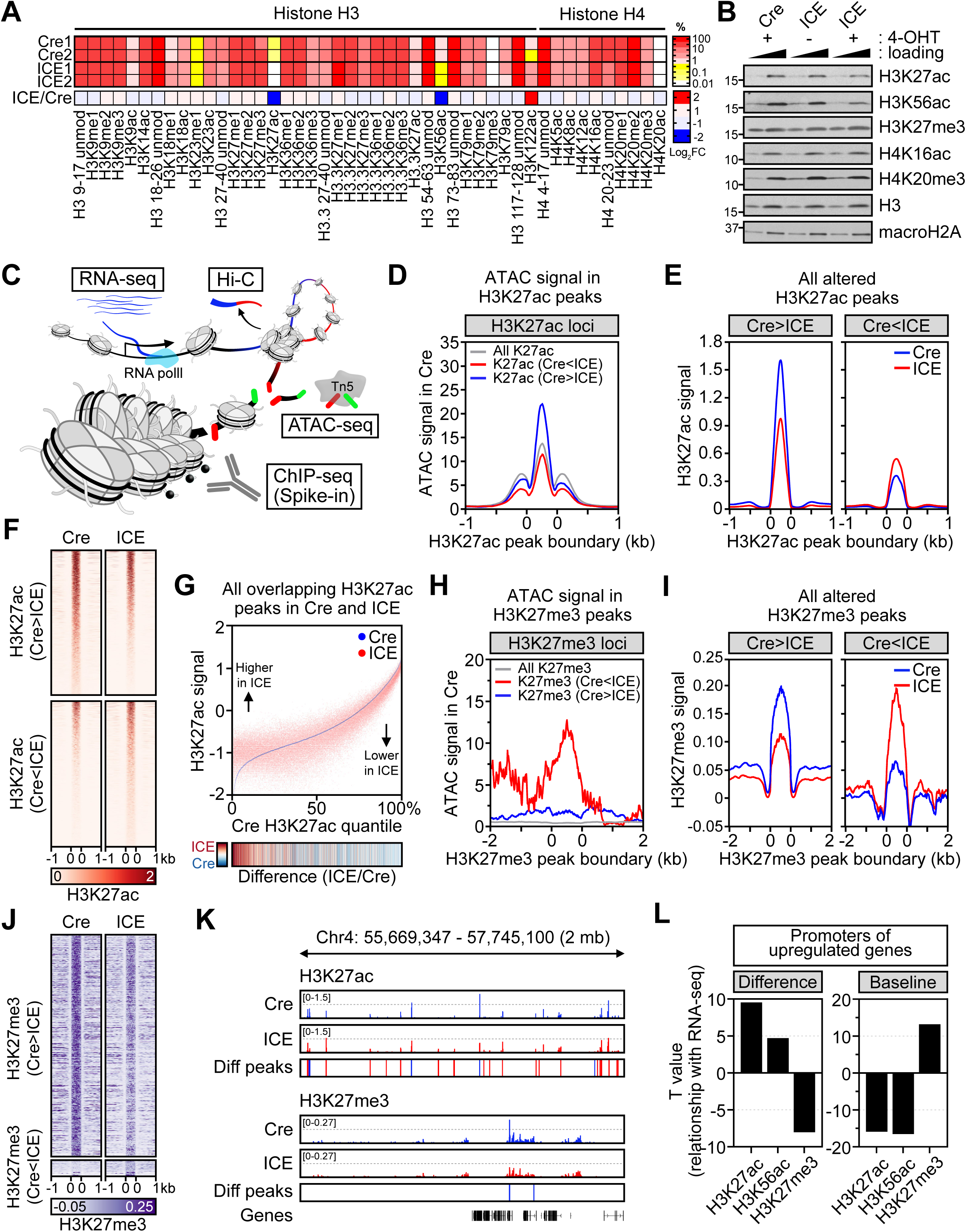
Smoothing of the Epigenetic Landscape in ICE Cells. (A) Quantitative mass spectrometry of histone H3 and H4 modifications in post-treated ICE cells; unmod, unmodified; me, methylation; ac, acetylation. (B) Western blotting for histone modifications in post-treated ICE cells. Histone H3, macroH2A and GAPDH serve as loading and internal controls. (C) Schematic of genome-wide analyses of the transcriptome (RNA-seq) and epigenome (ATAC-seq, ChIP-seq and Hi-C). (D) ATAC signal in H3K27ac peaks. (E and F) Aggregation plots and heatmaps of H3K27ac signal in H3K27ac changed regions (padj < 0.01). (G) Genome-wide changes of H3K27ac in ICE cells compared to Cre controls. Heatmap of ICE/Cre. (H) ATAC signal in H3K27me3 peaks. (I and J) Aggregation plots and heatmaps of H3K27me3 signal in H3K27me3 changed regions (padj < 0.01). (K) Representative region of chromosome 4 showing a smoothening of H3K27ac and H3K27me3 in post-treated ICE cells. Regions with changed peaks were marked in red (Cre<ICE) or blue (Cre>ICE). Diff peaks = ICE – Cre. (L) Correlation between histone modifications and mRNA levels. Difference, Cre vs. ICE; Baseline, signal in Cre.

Quantitative chromatin immunoprecipitation of Cre and ICE samples spiked with *Drosophila* S2+ cells (ChIP-Rx, ChIP-seq with reference exogenous genome) (Orlando et al., 2014) confirmed there was reduced chromatin-bound H3K27ac and H3K56ac in the ICE cells relative to Cre controls (2% and 5%, respectively) (**Figure S2A and Table S2**).

To map these changes with greater granularity (**Figure 2C**) we performed ChIP-seq for H3K27ac, H3K56ac and H3K27me3 and combined it with Assay for Transposase-Accessible Chromatin using sequencing (ATAC-seq) to map chromatin accessibility. Spearman correlations of ChIP-seq and ATAC-seq data showed the expected correlations (**Figure S2B**). At canonical I-*Ppo*I cut sites, I-*Ppo*I cutting had no effect on H3K27ac, H3K56ac, H3K27me3 or transcription (**Figure S2C and S2D**). Across the entire genome, the most significant change was H3K27ac, consistent with the bulk analysis of chromatin-bound and soluble H3 by mass spectrometry (**Figure S2E and Table S3**).

H3K27ac marks are generally associated with open chromatin, transcriptional activation, and cell type-specific enhancer activity (Heinz et al., 2015; Klemm et al., 2019). Consistent with this, genomic regions that lost H3K27ac in the treated ICE cells were more accessible, whereas regions that gained H3K27ac were less accessible (**Figure 2D**). An analysis of the aggregated H3K27ac signals that changed in the ICE cells showed that loci with high peak intensities tended to lose signal and vice versa, consistent with a smoothening of the H3K27ac landscape (**Figure 2E**, **2F**, **S2F**). Similar changes were seen with H3K56ac signals at the same regions and surrounding ATAC peaks (**Figure S2G and S2H**). A smoothening was also seen when assessing H3K27ac peaks genome-wide; the ICE:Cre ratio of H3K27ac was inversely correlated with the basal H3K27ac signal (**Figure 2G**, **S2I**, **S2J and S2K**).

Genomic regions that gained H3K27me3 in the ICE cells were also more accessible while the regions that lost H3K27me3 were less accessible, consistent with a loss of H3K27me3 from compact chromatin (**Figure 2H**). Similar to the behavior of H3K27ac, loci that had relatively high H3K27me3 peak intensities genome-wide (ICE vs. Cre) tended to lose the signal and vice versa, representing a smoothening of the H3K27me3 landscape (**Figure 2I and 2J**). At the mega-base scale, a smoothening of H3K27ac and H3K27me3 landscapes was even more evident, with a decrease in the high intensity peaks and an increase in the low intensity peaks (**Figure 2K**).

Because they target the same lysine, the acetylation and methylation patterns of H3K27 are often negatively correlated (Heinz et al., 2015). This was seen in the promoters of epigenetically aged ICE cells: upregulated genes gained both H3K27ac and H3K56ac but lost H3K27me3, with the promoters of downregulated genes showing the opposite (**Figure 2L and S2L**).

During development, cell identity is established and maintained, often for decades, by the polycomb repressive complex and H3K27me3 marks that silence loci that are used on other cells to specify a different cell type (Bernstein et al., 2006; Boyer et al., 2006; Lee et al., 2006).

Gene promoters that were upregulated in ICE cells tended to be enriched with H3K27me3 and depleted of H3K27ac at the basal level (**Figure 2L**), a correlation that is indicative of a decline in polycomb repression (Boyer et al., 2006; Lee et al., 2006). Together, these data indicate that DSB repair causes accessible chromatin associated with H3K27ac to lose this mark and regions of compact chromatin associated with H3K27me3 to lose polycomb suppression, consistent with a smoothening of the epigenetic landscape.

### Erosion of the Epigenetic Landscape Disrupts Developmental Genes

The increase in transcription in H3K27me3-marked promoters in the treated ICE cells (**Figure 2L**) indicated that their identity as fibroblasts might have been compromised. Gene Ontology (GO) analysis of the gene set that showed significant increases in H3K27ac, H3K56ac and decreases in H3K27me3 indicated that this was indeed the case. Within the top 20 processes, half of them were involved in developmental processes, including pattern specification, organ identity, tissue and organ development (**Figure 3A and Table S4**). This was further supported by GO analysis of genes that were upregulated in ICE cells at the mRNA level: 15 out of 20 were involved in developmental processes, including organ morphogenesis, limb morphogenesis, and the epithelial to mesenchymal transition (**Figure 3B and TableS6**). Consistent with RCM being an ancient stress response, loci that experienced a decrease in H3K27ac were predominately involved in stress responses, chromatin structure, metabolism, cellular component organization, nucleobase synthesis, and DNA repair (**Figure S3A and Table S4**). Genes involved in the metabolism of nitrogen and aromatic compounds were also enriched.

**Figure 3.**
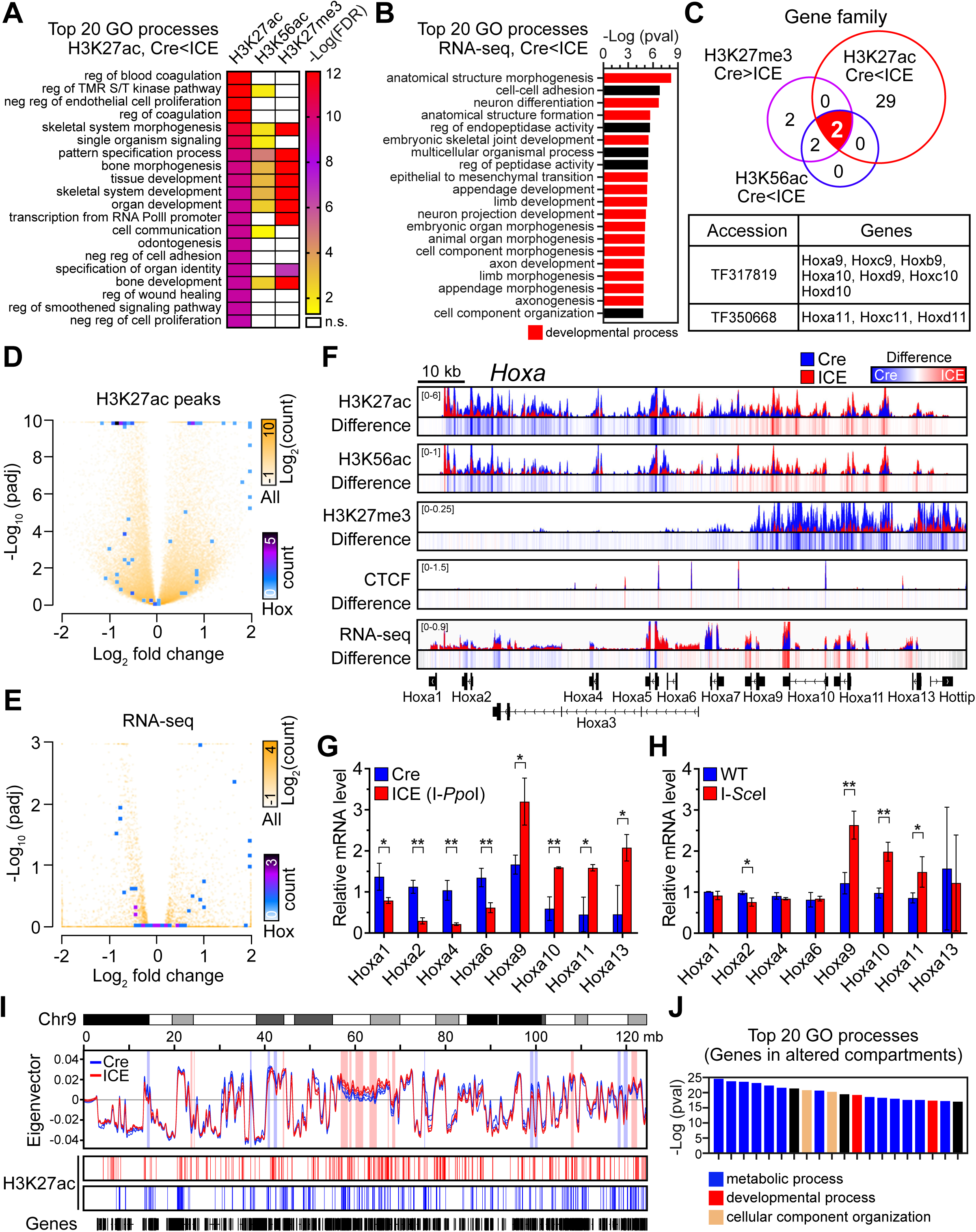
Dysregulation of Developmental Pathways in Epigenetically Aged ICE Cells. (A) Gene Ontology analysis of histone ChIP-seq data ordered by top 20 processes enriched in H3K27ac-increased regions (padj < 0.01). (B) Gene Ontology analysis of upregulated genes in RNA-seq data (padj < 0.05, FC > 1.5). (C) TreeFam analysis of gene families with overlapping regions with histone modification changes (padj < 0.01) in ICE cells. Two gene families with triple overlap were homeobox (*Hox*) genes. (D and E) Volcano plot of H3K27ac peaks and RNA-seq. All genes and *Hox* genes shown white to yellow and blue to purple, respectively. (F) ChIP-seq track of histone modifications and mRNA levels across the 120 kb *Hoxa* locus of post-treated ICE cells. (G and H) qPCR analysis of *Hoxa* genes in post-treated cells cut with either I-*Ppo*I or I-*Sce*I homing endonucleases. Two-tailed Student’s *t* test. (I) Eigenvector values for chromosome 9. Differentially compartmentalized regions were highlighted in red (Cre<ICE) or blue (Cre>ICE). Regions with altered H3K27ac peaks were marked in red (Cre<ICE) or blue (Cre>ICE). (J) Gene Ontology analysis of genes in differentially compartmentalized regions. Data are mean (n=3) ± SD. *p < 0.05; **p < 0.01.

To gain insights into the types of developmental processes affected in the ICE cells, the intersection of the ChIP-seq datasets in **Figure 3A** was cross-referenced with the TreeFam database, which provides orthology and paralogy predictions of gene families (Li et al., 2006; Ruan et al., 2008) (**Figure 3C**). At the intersection of all three data sets were two gene families, both comprised of homeobox (*Hox*) genes. These genes encode a conserved set of developmental transcription factors that specify body plan and the head-tail axis during embryogenesis and are found in gene clusters. In the epigenetically aged ICE cells, all of the *Hox* gene clusters (Hoxa-d) had significant alterations in peaks of H3K27ac, H3K56ac and H3K27me3 (**Figure 3D**, **3F**, **S3B and S3D-F**) coincident with changes in mRNA levels (**Figure 3E and S3C**).

Genes in the *Hoxa* locus were of particular interest with regards RCM because they are known to be regulated by SIRT1 and mediate stress and DNA repair responses (Oberdoerffer et al., 2008; Schworer et al., 2016; Singh et al., 2013). Across the *Hoxa* locus, there were clusters of genes that responded similarly. From *Hoxa1* to *Hoxa6*, levels of H3K27ac and H3K56ac decreased in ICE cells, while from *Hoxa9* to *Hoxa13* they increased, coincident with opposing changes in H3K27me3 and corresponding changes in mRNA levels (**Figure 3F and G**). These data add further evidence that repeated cycles of DSB repair alter chromatin in a way that erodes boundaries and smoothens the epigenetic landscape.

To test if the effects of DSB repair were depended on the I-*Ppo*I cut site, we isolated MEFs from a mouse strain with an inducible homing endonuclease from budding yeast called I-*Sce*I, which cuts at ∼18 cryptic (non-canonical) sites in the mouse genome, far from the vicinity of I-*Ppo*I sites (Chiarle et al., 2011). Paralleling the effects of I-*Ppo*I on post-recovery gene expression, I-*Sce*I altered mRNA levels of genes in the *Hoxa* and *HIST* cluster. Thus, the effect of DSBs on Hoxa expression does not depend on where the DNA breaks occur (**Figure 3H and S4C**).

Global histone expression has been shown in yeast to decline with age, the repletion of which extends yeast lifespan. This and other findings indicate that histone loss is a cause of aging in yeast and possibly mammals (Feser et al., 2010; Hu et al., 2014). Indeed, the locus that showed one of the most significant and robust epigenetic change in ICE cells was histone cluster 1 (*Hist1*), which encodes canonical histones H1, H2A, H2B, H3 and H4. Though there was no increase in senescence of the ICE cells at this time point, similar decreases in histone expression have also been seen in senescent cells, and in cells from aged rodents, such as muscle satellite cells (Liu et al., 2013). Across the *Hist1* locus, there was a global decrease in H3K27ac and H3K56ac along with decreases in mRNA and protein levels (**Figure S4A-S4D**). Decreases in H3 and H4 mRNAs were also seen in post-treated I-*Sce*I cells (**Figure S4C**). Reduced histone levels could have been due to changes in the cell cycle, such as a slower S-phase, but no difference in cell cycle profiles were detected using a variety of approaches, including FACS and incorporation of the thymidine analogue 5-ethynyl-2’-deoxyuridine (EdU) into replicating DNA (**Figure S4E-S4F**; Hayano et al, co-submitted manuscript).

### DSB Repair Alters Chromatin Compartmentalization

Transcription factor networks are essential for maintaining cellular function and identity. Transcription factor (TF) binding enrichment analysis of loci with reduced levels of H3K27ac and H3K27me3 identified of a total of 18 potentially relevant TF binding sites, 6 of which are known to be involved in DNA repair or development (**Figure S4G and Table S5**). One of these was CTCF, a protein that binds to DNA and multimerizes to facilitate DNA repair and the formation of topologically associated domains (TAD) (Dixon et al., 2012; Lang et al., 2017). CTCF had highly significant p values of 2.2 x 10^−130^ and 2.4 x 10^−5^ for H3K27ac and H3K27me, respectively (**Figure S4G-S4H**). Although there was no change in CTCF protein levels or CTCF binding globally, among CTCF peaks that did change (p<0.01), there was a clear smoothening of the CTCF landscape in post-treated ICE cells (**Figure S4I-S4K**). CTCF binding sites were preferentially associated with decreased H3K27ac signals compared to non-CTCF binding sites (p < 10^−199^), consistent with the TF binding site enrichment analysis (**Figure S4L**). Recovered ICE cells had ∼35% more CTCF foci than matched Cre cells. Together these data indicating that CTCF multimers may have been disrupted (**Figure S4M**).

To detect changes in chromatin architecture, proximity ligation (Hi-C) experiments were performed on Cre and ICE cells. Generally, TADs are robust structures that show little variation, even across different cell types (Dixon et al., 2015; Dixon et al., 2012; Rao et al., 2014) but chromatin compartmentalization, represented as a Hi-C eigenvector, is a property that emerges independently of TADs (Nora et al., 2017; Rao et al., 2017). At the level of TADs, there were no significant changes in post-treated ICE cells (**Figure S3G**) but there were highly reproducible differences in compartmentalization that were also enriched for H3K27ac changes. GO analysis revealed that genes within these compartments were significantly enriched for developmental processes, metabolism, and neurogenesis (**Figure 3I and 3J**). To our knowledge, DNA damage-induced changes in chromatin compartmentalization at sites unrelated to the site of DNA damage, what we refer to as “distal chromatin scars,” have not been previously described.

### Epigenetically Aged Cells Lose the Ability to Maintain Cellular Identity

The smoothening of the epigenetic landscape across the genome and the fact that the most prominent epigenetic and transcriptional changes were in genes involved in developmental pathways and cellular differentiation, prompted us to test if the identity of the ICE fibroblasts has been impacted. In mammals, cellular identity is established prenatally and maintained by chromatin regions called super-enhancers (SE), large clusters of smaller enhancers that are dense with enhancer-associated proteins, H3K27ac marks (Hnisz et al., 2013; Whyte et al., 2013) and broad H3K4me3 signals (Benayoun et al., 2014). A comparison of altered H3K27ac peak signals and the super-enhancer database, dbSUPER (Khan and Zhang, 2016), revealed that the most significant loss of H3K27ac marks in the ICE cells occurred at MEF SEs, consistent with ICE cells losing their identity (**Figure 4A and 4B**). There was also a clear loss of the H3K27ac signal at the top 5% of H3K4me3 broad domains (**Figure S5A and S5B**) and a decrease in H3K27ac marks and mRNA levels of canonical genes that specify fibroblast lineages, including *Col1A1* and *Thy1* (**Figure S5C-S5F**). Similar changes were seen at *Chaf1a* and *Chaf1b*, two histone chaperone genes that maintain somatic cell identity (Cheloufi et al., 2015) (**Figure S5G and S5H**).

**Figure 4.**
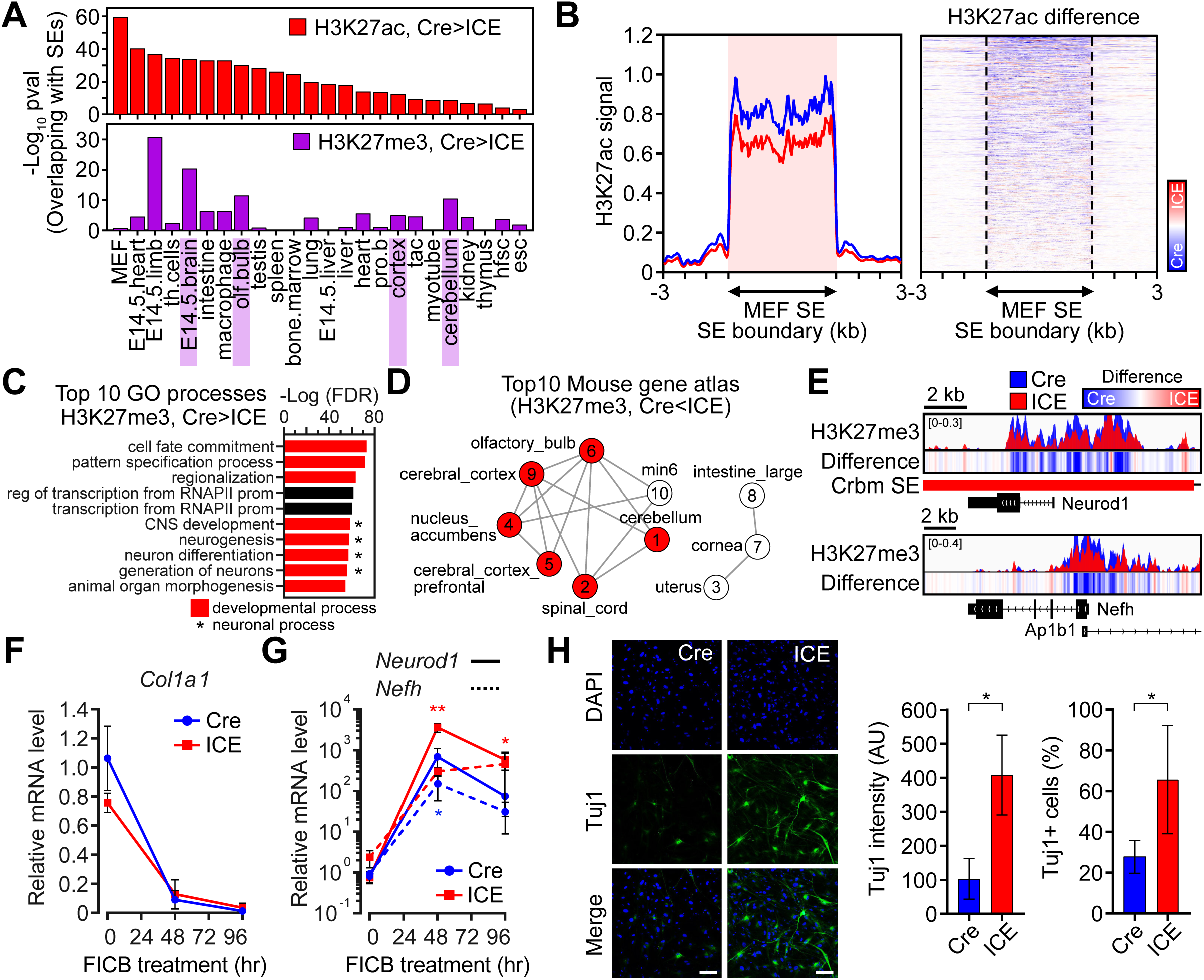
Loss of Cellular Identity in Epigenetically Aged ICE Cells. (A) Super-enhancers (SEs) in different cell types that overlap with regions with reduced H3K27ac and H3K27me3 signals (Cre>ICE in MEFs). (B) Aggregation plots and heatmaps of H3K27ac signals in MEF SE regions. (C) Gene Ontology analysis of H3K27me3 decreased regions (padj < 0.05). Red represents developmental processes. *neuronal processes. (D) Mouse tissue types of transcriptional profiles that overlap decreased H3K27me3 regions (padj < 0.05) in epigenetically aged ICE cells. Red represents neuronal tissues. Numbers indicate ranks. (E) ChIP-seq track of representative neuronal marker genes, *Neurod1* and *Nefh*. Diff peaks = ICE – Cre. (F and G) Time-course analysis of mRNA levels of *Col1A1* (a fibroblast marker), *Neurod1* and *Nefh* (neuronal markers) during neuronal reprogramming. Two-way ANOVA-Bonferroni. (H) Immunostaining and quantification of neuronal marker Tuj1 after 8 d of reprogramming. DNA stained with DAPI. Scale bar, 100 µm. Two-tailed Student’s *t* test. Data are mean (n≥3) ± SD. *p < 0.05; **p < 0.01.

The H3K27me3 mark maintains cellular identity by suppressing super-enhancers that specify other cell types (Adam et al., 2015; Tee and Reinberg, 2014). At cell type SEs, post-treated ICE cells had weaker H3K27me3 signals, three out of four of which were SEs that dictate neuronal cell types in the embryonic brain, olfactory bulb and cerebellum (**Figure 4A**). An assessment of GO processes or tissue types of transcriptional profiles that overlap genes with decreased H3K27me3 revealed that 4 and 6 out of the top 10 were involved in neuronal processes and tissue types including cerebral cortex, spinal cord, and cerebellum (**Figure 4C**, **4D and S5I**). H3K27me3 signals were lower across promoter regions of genes that specify neuronal fate, including the *Neurod1* gene, which lies within a cerebellum SE, and the neurofilament gene *Nefh,* which maintains neuronal caliber (**Figure 4E**).

These findings indicated that epigenetically aged MEFs might have shifted away from a fibroblast lineage and more towards a neuronal landscape. If so, they should be primed to differentiate into neuronal cell types. To test this, Cre and ICE post-treated MEFs were subjected to a standard 17-day neuronal reprogramming protocol that uses small molecules to induce neuronal genes and inactivate fibroblast genes such as *Col1A1* (Li et al., 2015) (**Figure 4F**, **4G**, **S6A and S6B**). During reprogramming, *Neurod1* and *Nefh* were 8-15-fold more easily derepressed in ICE MEFs than in Cre control cells (**Figure 4G and S6C**). Compared to the Cre controls, 2.5-fold more ICE-derived neurons were created along with 4-fold higher levels of Tuj1, a canonical neuronal cell marker (**Figure 4H**).

The trans-differentiation of MEFs to adipocytes is another well-established system for evaluating cellular plasticity. Unlike neurons, which must be generated from embryonic or iPS cells, the fibroblasts of adult mice can be trans-differentiated into adipocytes, allowing us to test if ICE cells behaved like normal aged cells (Alexander et al., 1998). Prior to treatment, compared to Cre controls, ICE cells had significantly more H3K27ac in the enhancer region of the adipocyte master regulator *Pparg* (**Figure S6D**) and after seven days of treatment with adipogenic reprogramming factors, there was a significant increase in lipid droplets in ICE cells relative to Cre controls, paralleling the known effect of aging (**Figure S6E-S6F**).

### Gene expression changes in ICE cells resemble aging

Another robust prediction of the RCM hypothesis is that gene expression changes in cells post-DSB repair should resemble cells taken from older animals. To test this, we performed RNA-seq on adult ear fibroblasts from mice aged 3, 24, and 30 months, and compared them to Cre and ICE MEFs. Of the top 20 pathways elevated in the fibroblasts from 24 and 30 month-old mice, 14 were in developmental or cellular differentiation pathways (**Figure S7A, Table S7**), all of which were also elevated in the epigenetically aged ICE cells. Of the top 20 GO biological processes altered in ICE cells, 15 of them were developmental or cellular differentiation processes (**Figure 3B and Table S7**). Compared to the 3-month samples, 19 of the 20 changes in ICE MEFs were also significantly enriched in fibroblasts from old mice, a high degree of overlap between ICE fibroblasts aged *in vitro* and wildtype fibroblasts aged *in vivo* (**Figure 5A).** At the individual gene level, 275 mRNAs were significantly different in both the epigenetically-aged ICE cells and fibroblasts from old mice, the majority of which changed in the same direction (**Figure 5B and 5C**). The cells from old mice also had dysregulation of *Hoxa*, histone genes, *Lmnb1* and *Chaf1a*, paralleling what we observed in the ICE cells post-treatment (**Figure 5D**, **5E and S7B**). Changes in developmental processes and similar patterns of mRNA dysregulation were also seen in human fibroblasts from older individuals (Mertens et al., 2015) (**Figure 5F**, **5G and S7C**).

**Figure 5.**
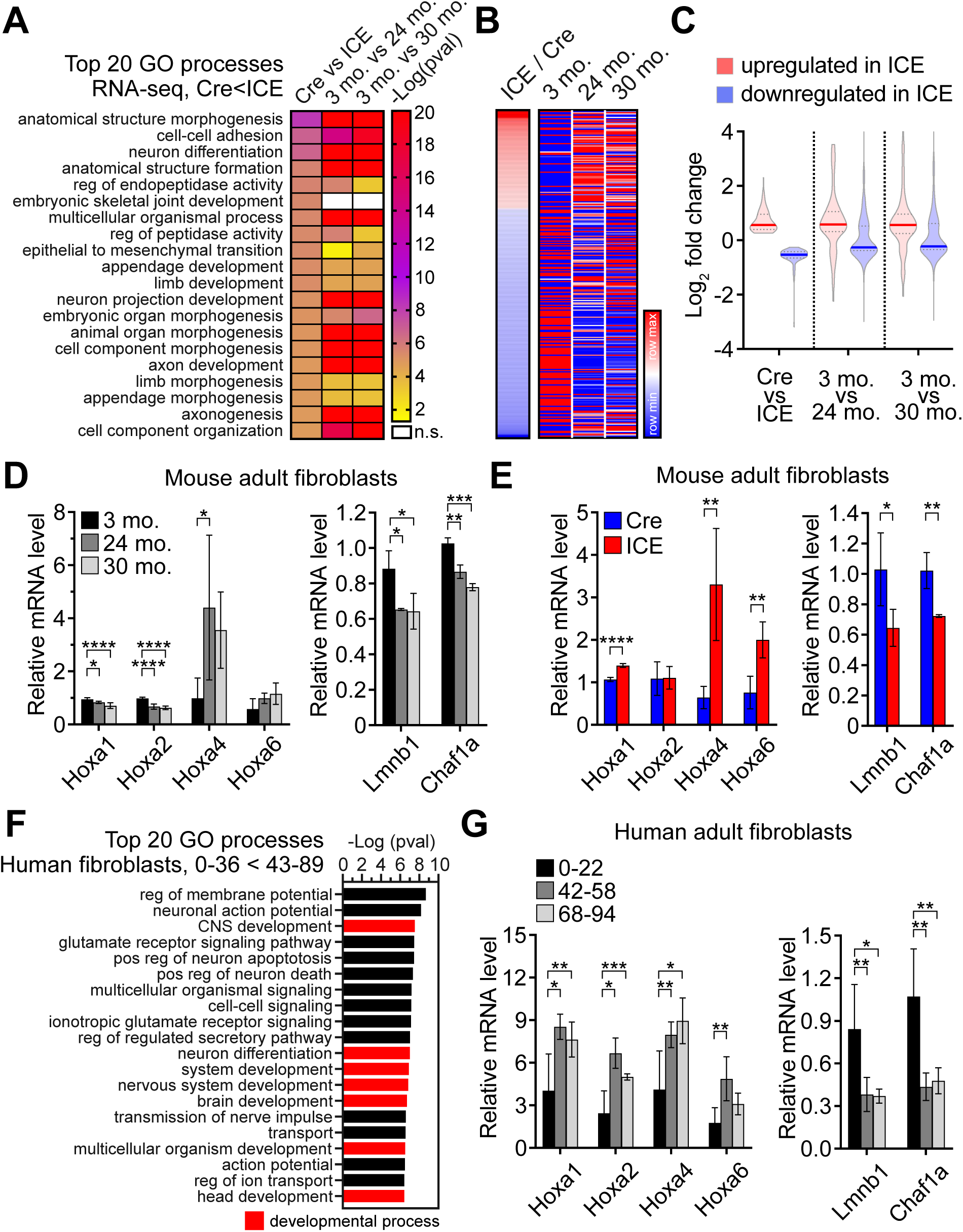
Epigenetically Aged ICE Cells Mimic Cells from Old Mice. (A) Gene Ontology analysis of up-regulated genes (padj < 0.05, FC > 1.5) in RNA-seq data from epigenetically aged ICE cells and fibroblasts from young and old mice. (B) Heatmaps of significantly altered mRNA levels in ICE cells compared to genes from cells from young and old mice (p < 0.01, both in ICE and old cells). (C) Mean log_2_ fold change of mRNA up- or down-regulated in ICE cells compared to cells from young and old mice. (D and E) mRNA levels of *Hoxa*, *Lmnb1*, and *Chaf1a* in adult fibroblasts from 3, 24, and 30 month-old mice and 6 month-old ICE mice. One-way ANOVA-Bonferroni (left), two-tailed Student’s *t* test (right). (F) Gene Ontology analysis of RNA-seq data from human fibroblasts from young and old individuals. (G) mRNA levels of genes in (D) in human adult fibroblasts of different ages. One-way ANOVA-Bonferroni. Data are mean (n≥3) ± SD. *p < 0.05; **p < 0.01; ***p< 0.001; **** p < 0.0001.

### Cellular Identity Changes in ICE Mice

A hallmark of mammalian aging is a switch in cellular identity that resembles the epithelial to mesenchymal transition (EMT) (Brack et al., 2007; Roeder et al., 2015; Schneider et al., 2017). Processes involved in the EMT were significantly altered in the prematurely-aged ICE and normally-aged fibroblasts (**Figure 5A**). In ICE cells, regions that gained H3K27ac or lost H3K27me3 marks significantly overlapped with those involved in the establishment and maintenance of cellular identity in muscle and kidney, two tissues known to lose their identity over time (Brack et al., 2007; Roeder et al., 2015) (**Figure 6A**).

**Figure 6.**
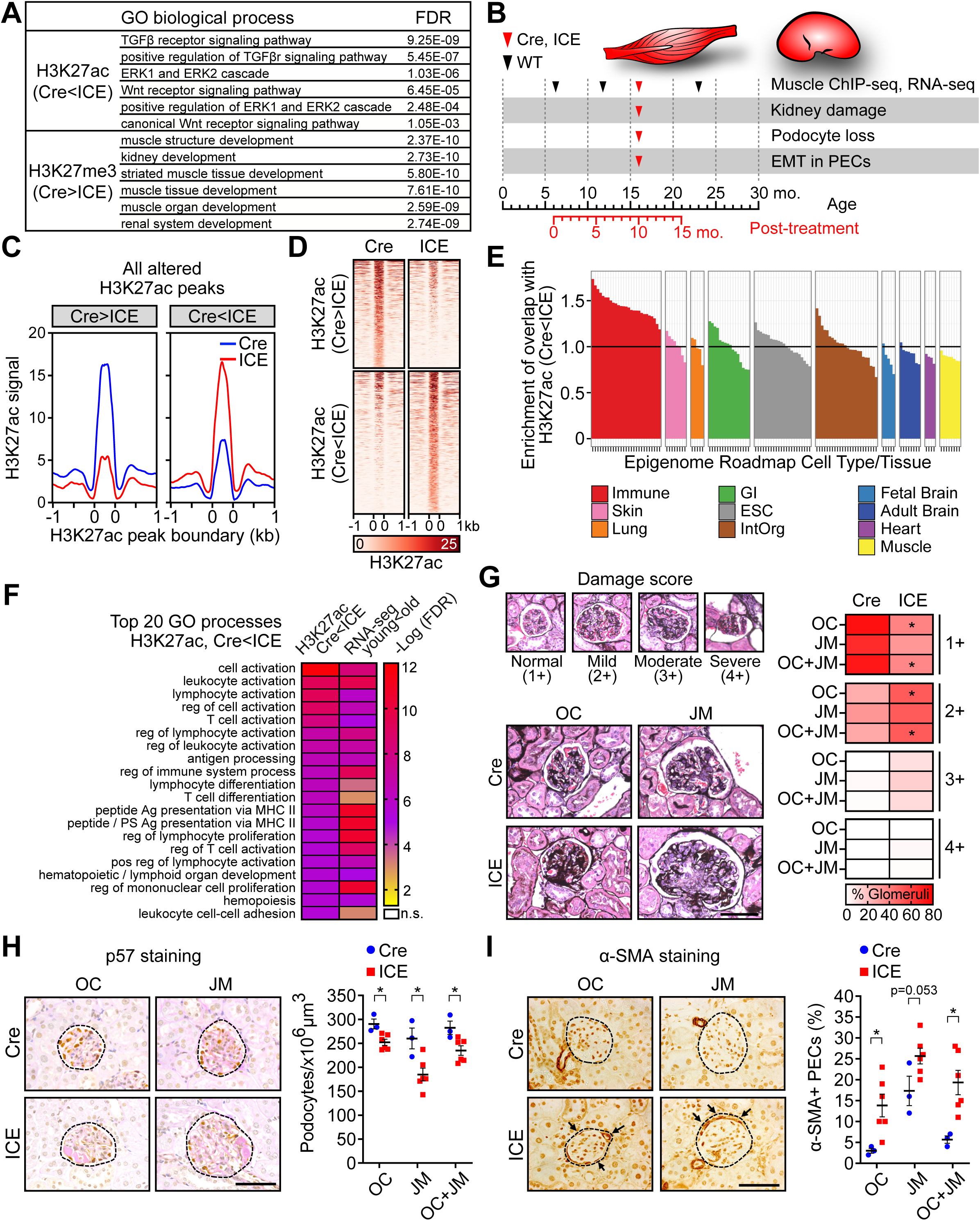
Tissues of ICE Mice Lose Cellular Identity, Mimicking Old Wildtype Mice. (A) Gene Ontology processes involved in muscle and kidney development that were significantly enriched for increased H3K27ac peaks (padj < 0.01) and decreased H3K27me3 peaks (padj < 0.01). (B) Timeline of tissue analyses in ICE and wildtype mice. (C, D) Aggregation plot and heatmaps of H3K27ac changed regions (p < 0.01) in skeletal muscle. (E) Comparison of H3K27ac increased regions (p < 0.01) to epigenome roadmap data from different human tissue types. (F) Gene Ontology comparison of H3K27ac increased regions in epigenetically aged ICE mice (16 mo.) (p < 0.01) to RNA-seq data from skeletal muscle from old wildtype mice (24 mo.) (padj < 0.05). (G) Representative images and average damage scores (1+ normal – 4+ global scarring) of glomeruli of 16 month-old ICE mice. OC, outer cortex; JM, juxtamedullary glomeruli. Two-tailed Student’s *t* test. (H) Representative images of p57 (podocyte) and Periodic acid-Schiff staining and podocyte density of 16 month-old ICE mice. Circles with broken line indicate glomeruli. Scale bar, 50 µm. Two-tailed Student’s *t* test. (I) Representative images and fraction of α-SMA-positive cells in parietal epithelial cells (PEC) along Bowman’s capsule (arrows) of 16 month-old ICE mice showing an epithelial to mesenchymal transition (EMT). Circles with broken line indicate glomeruli. Scale bar, 50 µm. Two-tailed Student’s *t* test. Data are mean (n≥3) ± SEM. *p < 0.05.

**Figure 7.**
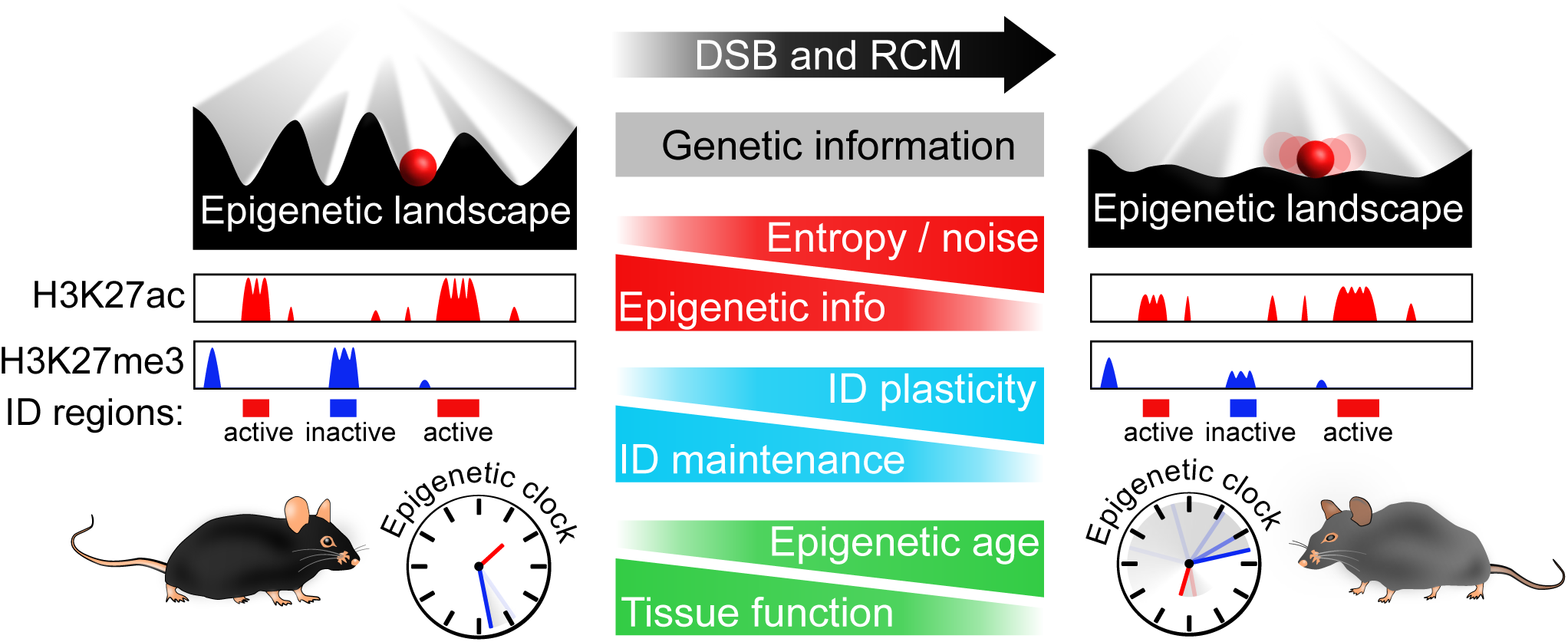
Model for the Loss of Epigenetic Information during Aging. During development, cell types are specified by the establishment of specific transcriptional networks and chromatin landscapes, such that cells land in valleys depicted by the Waddington epigenetic landscape. Repeated disruptions to the epigenome––such as DSBs, which cause dual-function chromatin modifying DNA repair factors to relocalize––cause the landscape to erode and the epigenetic clock to advance. Cells eventually move closer to nearby valleys as they lose their identity and specific functions, culminating in aging.

To test whether DSBs also accelerate the loss of cell identity *in vivo*, we induced I-*Ppo*I for three weeks in the entire body of 4-6 month-old ICE mice by feeding them with tamoxifen (360 mg/kg) for three weeks. Ten months later, mice were sacrificed and RNA-seq and ChIP-seq were performed on skeletal muscle (**Figure 6B**). In the gastrocnemius muscle there was an increase in H3K27ac low intensity peaks and a decrease in the high intensity peaks (**Table S8 and S9**), paralleling what we had observed in the ICE cells, though more subtle and at a lower frequency than ICE MEFs (1 vs. 20%) (**Figure 6C**, **6D and S7D**).

To assess if the muscle of ICE mice had taken on a signature of another cell type, we compared the dataset of regions with altered H3K27ac signals to the epigenome roadmap, a consortium of human epigenomic data from different cell types and tissues (Roadmap Epigenomics et al., 2015). Regions with lower H3K27ac (p < 0.01) in Cre vs. ICE showed the strongest enrichment for muscle tissue signatures (p = 9.0×10^−8^) while regions with higher H3K27ac showed an enrichment for immune cell enhancers (p = 9.3×10^−28^) (**Figure 6E and S7E**). Of the regions that gained H3K27ac, there was overlap with super-enhancer regions from immune cell types and with regulatory regions involved in developmental processes and immune cell activation (**Figure 6F**, **S7F and S7G**).

Of the top 20 processes in ICE muscle, all 20 were also elevated in old wildtype mice, indicating the ICE mice closely paralleled changes seen in normal aging (**Figure 6F**). Regions with decreased H3K27ac signals overlapped with processes that included metabolic processes, response to stress, and cellular component organization, consistent with ICE cells (**Figure S7H**). Because there was no evidence of immune cell infiltration into the muscle of ICE mice (**Figure S7I**) and ChIP-seq data is not influenced by a small portion of infiltrated cells, we infer that the muscle tissue in treated ICE mice had shifted more towards an immune cell. The observed increased overlap of inflammation-related gene expression patterns and chromatin changes in old mice is consistent with a recent report that examined a variety of tissues, including muscle (Benayoun et al., 2019).

Besides muscle, the other tissue that significantly overlapped with changes to histone marks in the ICE MEF was kidney, an organ often used as a model for EMT during aging (**Figure 6A**). Two well established age-related changes in kidney are an increase in damage of the glomerular basement membrane and loss of podocytes that line the glomerular capillary. As in old wildtype mice (Roeder et al., 2017), the kidneys of treated ICE mice had fewer tissue samples that scored as normal (1+) compared to Cre controls, and a higher percent of those scored as damaged (2+ and 3+) for both the outer cortex (OC) and juxtamedullary (JM) glomerulus, along with a greater loss of podocytes (**Figure 6G-H**). A well-known marker for EMT is the alpha isoform of smooth muscle actin (αSMA). ICE mice had increased αSMA staining in both the OC and JM relative to Cre controls, paralleling changes seen during normal aging (**Figure 6I**) (Roeder et al., 2017). Together with the *in vivo* data in the co-submitted manuscript (Hayano et al.), these analyses indicate that the induction of non-mutagenic DSBs accelerates the DNA methylation clock and induces many of the same changes to chromatin, gene expression, and cellular identity that occur in normal mouse aging.

## DISCUSSION

The RCM hypothesis of aging predicts that the reorganization of chromatin to address threats to cell survival, such as a broken chromosome, introduces epigenetic noise that causes cells to lose their identity and become dysfunctional. Indeed, this hypothesis is well established in budding yeast. But whether or not it is true for mammals is debated. The ICE system allowed us to introduce epigenetic noise into cultured cells and mice and test the prediction of the RCM hypothesis: that repeated non-mutagenic DSB repair events should accelerate the epigenetic clock and induce both chromatin and physiological changes that mimic aging. Indeed, in the accompanying manuscript, we show that the introduction of epigenomic noise in mice closely replicates an aged phenotype at the molecular and physiological levels, including an acceleration of the epigenetic clock (Hayano et al, co-submitted manuscript).

In this part of the two-part study, we sought to understand the process at a molecular level, specifically whether DSBs leave an epigenetic scar and what impact that has on chromatin architecture, gene expression, and cell identity. We present evidence that the response to DSBs changes the compartmentalization of chromatin and introduces transcriptional and epigenetic noise that closely mimics what happens during normal aging, including hallmark changes to the histone modifications, gene expression, and DNA methylation patterns. One of the most notable changes *in vitro* and *in vivo* was a loss of epigenetic signatures at super-enhancers that dictate cellular identity. These changes were part of an overall smoothening of the epigenetic landscape with decreased signal to noise ratios that did not appreciably affect the cell cycle but set off a chain of events culminating in cellular senescence. Consistent with this, senescent cells were recently reported to also undergo a smoothing of the epigenetic landscape (De Cecco et al., 2013).

One of the more notable findings in the past decade is that mammalian aging is accompanied by a deregulation of developmental pathways, including Wnt, TGFβ, and ERK, leading to a loss of cellular identity (Liu et al., 2007; Schworer et al., 2016). In tissues such as muscle and kidney, this has been well established (Brack et al., 2007; Roeder et al., 2017). In this study, we present evidence that this process is initiated, at least in part, by the reorganization of chromatin in response to DSBs and a smoothening of the epigenetic landscape. We speculate that changes to histone modifications in ICE cells precede changes in CTCF binding and TADs, but ultimately they are disrupted due to a positive feedback loop driven perhaps by a loss of histones and increased susceptibility to DNA damage, causing cells to eventually senesce, even if they are non-dividing (Criscione et al., 2016; Zirkel et al., 2018). In the context of the Waddington landscape, our data suggests that cell types in adjacent valleys, having arisen from the same lineage during development, are more likely to interconvert during aging, such as fibroblasts to neurons, which are both derived from the neuro-ectodermal lineage, and muscle to immune cells, which are both derived from the mesoderm (Benayoun et al., 2019).

We do not know why DSBs preferentially disrupt developmental pathways, but one explanation is that other classes of genes regulated by transient transcriptional networks are more easily reset to their original state after DSB repair than developmental genes, which are under multiple layers of regulation. Based on striking parallels with yeast aging, the coordination of DSB signaling with cellular identity appears to be an ancient mechanism, one that may have evolved to temporarily invoke DNA repair mechanisms in embryonic cells, which must deal with an abundance of potentially lethal DSBs that occur as a result of rapid DNA replication. This would explain why developmental regulators, such as Wnt and Hox genes, also mediate DSB repair (Feltes, 2019; Rubin et al., 2007; Zhao et al., 2018).

Together, the data in this study and the accompanying manuscript (Hayano et al, co-submitted manuscript) are consistent with the following model. During embryonic development, each cell type is endowed with an epigenetic landscape that allows cells to maintain cellular identity for decades, even while transcriptional networks adapt to an ever-changing environment. Potent disrupters of the epigenome, such as DSBs, temporarily disrupt chromatin architecture and recruit chromatin factors away from certain loci to assist with repair and relieve repression of stress response and DNA repair genes. After repair, the epigenome is reset but not completely, leading to progressive changes to the epigenetic marks and chromatin compartmentalization of the genome. Due to a loss of histones and the suppression of stress response genes, the genome becomes more susceptible to DSBs, creating a vicious cycle. In the parlance of Waddington, the youthful epigenetic landscape is eroded to the point where cells head towards other valleys, losing their identity in a process we have termed “exdifferentiation.” Ultimately, when erosion reaches a critical threshold and DNA damage signaling is constitutively activated, cells senesce or perhaps become tumorigenic if certain mutations are present (Fumagalli et al., 2014; Wang et al., 2019).

The linking of DSB signaling to mammalian aging raises a number of interesting questions. For example, how exactly do DSBs accelerate the DNA methylation clock? One possibility is that DNA methylation is altered at sites of DNA repair. Indeed, the DNA methyltransferases DNMT1 and DNMT3B are recruited to DSBs altering DNA methylation, and TET2-dependent enrichment of 5hmC at DNA damage sites is needed for genome stability (Kafer et al., 2016). Given that the DNA methylation sites in the clock did not overlap with the I-*Ppo*I cut sites in our system, it seems unlikely the DNA methylation changes are direct. We envisage that DNA methylation enzymes may be recruited away from distal loci, in the same way SIRT1 and HDAC1 are (Dobbin et al., 2013; Oberdoerffer et al., 2008). Because in this model it is not the site of damage but the reaction to it that causes aging, it would explain how random DNA damage can induce a predictable pattern of epigenetic and physiological changes over a lifetime leading to a common set of age-related diseases.

Our data also indicates that DSB signaling, even in the absence of actual DSBs, should be sufficient to drive aging. It will be interesting to assess the role of specific chromatin factors in the ICE model, such as SIRT1/SIRT6, PARP1, HDAC1, EZH2, DMNTs and TETs, in tissues and at the single cell level in aging, progeroid syndromes, and the ICE system. Transcription factors such as CTCF or FOXO, whose binding motifs were significantly enriched in altered H3K27ac regions in both ICE MEFs and old muscle, can direct these chromatin factors (**Table S5**).

For over 50 years, a loss of genetic information was considered the main cause of aging. Recent studies, including this one, however, indicate that aging may be caused in large part, by a loss of epigenetic information. The second law of thermodynamics states that the total entropy of an isolated system can never decrease over time and therefore all ordered systems eventually succumb to noise. But living things are exception because they are open systems. According to Shannon (Shannon, 1948), a viable solution is to have an “observer” who retains a copy of original information so, if it is lost to noise, it is later recoverable. If cells retain such information, it may be possible to restore youthful epigenomic landscapes in old cells to reverse many of the effects of aging on tissues and organs.

## Supporting information

Supplemental Information_Yang et al

## ACKNOWLEDGMENTS

We thank all members of the Sinclair laboratory for constructive comments on the project and manuscript, Paul F. Glenn, Edward Schulak and AFAR for generous support. We also thank to Roberto Chiarle (Boston Children’s Hospital) and Frederick Alt (Harvard Medical School) for sharing I-*Sce*I mice and Haeyoung Kim (Texas A&M University-Kingsville) and Peter Adams (Sanford Burnham Prebys Medical Discovery Institute) for technical advice. Research was supported by NIH/NIA (R01AG019719 and R37AG028730 to D.A.S.), the Glenn Foundation for Medical Research (to D.A.S.), NIH (5R01DK056799-10, 5R01DK056799-12, 1R01DK097598-01A1 to S.J.S.), National Research Foundation of Korea (2012R1A6A3A03040476 to J.-H.Y.), Human Frontier Science Program (LT000680/2014-L to M.H.), NIH T32 (T32AG023480 to D.L.V.) and Glenn/AFAR Research Grants for Junior Faculty (to A.R.P.).

## AUTHOR CONTRIBUTIONS

J.-H.Y., L.A.R. and D.A.S. initiated and designed the project. J.-H.Y. performed most experiments, analyzed data and wrote the manuscript. A.R.P., P.T.G., D.L.V., J.A., J.-H.Y., and C.S. analyzed ChIP-seq, RNA-seq and ATAC-seq data. E.L.S. analyzed WGS data. M.H. performed muscle ChIP-seq. D.L.V., J-H.Y., E.M.M., M. Blanchette, M. Bhakta, B.O., and R.E.G. performed and analyzed Hi-C. Z.D., C.X., B.A.G. and S.L.B. performed and analyzed MS. M.V.M. and V.N.G. performed RRBS and calculated epigenetic ages. J.W.P., M.C. and S.J.S. performed kidney staining and analysis. P.O. designed the I-*Ppo*I construct and integrated the construct into ES cells to generate I-*Ppo*I mouse. L.A.R. initiated ICE mouse study and provide advice and assistance throughout. D.A.S. supervised the project and wrote the manuscript.

## METHOD DETAILS

### Cell culture

Mouse Embryonic Fibroblast (MEF) cells were isolated from E13.5 mouse embryos. After dissecting out the uterus and yolk sac, fetuses were moved in a new dish containing sterile PBS. The liver, heart, head were removed and the remaining part was washed in sterile PBS to remove blood. Fetuses were minced in 0.25% trypsin-EDTA and incubated at 37°C for 30 min. Cells were washed and maintained with MEF growth medium (DMEM containing 20% FBS, 1% penicillin/streptomycin, 0.1 mM β-mercaptoethanol). For activation of ER (estrogen receptor)-fused Cre in MEFs, 0.5 μM 4-Hydroxytamoxifen (4-OHT) was treated for 24 h and medium was switched to one without 4-OHT to stop I-*Ppo*I-mediated DNA breaks. For activation of GR (glucocorticoid receptor)-fused I-*Sce*I, 10 μM triamcinolone acetonide (TA) were treated in DMEM containing charcoal stripped FBS for 2 days.

Mouse adult fibroblast cells were isolated from ears taken from 3, 24 and 30 month-old mice. 2 whole ears were washed with 70% EtOH and sterile PBS and minced in DMEM containing 0.14 Wunsch Units/ml Liberase TM and 1% penicillin/streptomycin. After incubation of minced tissues at 37°C for 45 min with shaking, cells were washed with medium twice and plated on collagen coated culture dishes.

Human adult fibroblast cells were obtained from Coriell Institute.

All cells were cultured in DMEM containing 20% FBS (Seradigm), 1% penicillin/streptomycin, 0.1 mM β-mercaptoethanol at 37°C, 3% O_2_ and 5% CO_2_ unless otherwise specified.

### ChIP-sequencing

MEF ChIP was done following the protocol described in (Yang et al., 2011) with minor modifications. 1/4 number of *Drosophila* S2R+ cells relative to mouse cells were added as a spiked-in control and combined cells were treated as a single sample during the rest of the procedures. Cells were cross-linked with 1% formaldehyde at RT for 10 min and glycine was added to final concentration 0.125 M for 5 min to quench crosslinking. Fixed cells were washed with PBS and nuclei were isolated using Lysis buffer A (10 mM Tris-HCl pH 7.5, 10 mM KCl, 5 mM MgCl2, 0.5% NP40, protease inhibitor cocktail). Nuclei were resuspended in SDS lysis buffer (50 mM Tris-HCl pH 7.9, 10 mM EDTA, 0.5% SDS, protease inhibitor cocktail). Chromatin was sheared using Covaris E210 Ultrasonicator (duty cycle:5%, intensity:4, cycle/burst:200, time:15-20 min) to generate fragmented chromatin ranging between 200 and 1,000 bp. After centrifugation, sonicated chromatin solution was 5 fold-diluted with ChIP dilution buffer (12.5 mM Tris-HCl pH 7.9, 187.5 mM NaCl, 1.25% Triton X-100, protease inhibitor cocktail).

Antibodies and magnetic beads were added to diluted chromatin solutions and immunoprecipitation were performed at 4°C overnight with rotation. Immunocomplexes were washed with Low salt wash Buffer (0.1% SDS, 1% Triton X-100, 2 mM EDTA, 20 mM Tris-HCl pH 8.1, 150 mM NaCl), High salt wash buffer (0.1% SDS, 1% Triton X-100, 2 mM EDTA, 20 mM Tris-HCl pH 8.1, 500 mM NaCl), LiCl wash buffer (0.25 M LiCl, 1% NP40, 1% deoxycholate, 1 mM EDTA, 10 mM Tris-HCl pH 8.1), and TE (10 mM Tris-HCl pH 8.0, 1 mM EDTA).

Immunocomplexes were eluted in elution buffer (1% SDS, 0.1 M NaHCO3) at RT for 30min with rotation, and RNaseA (final concentration of 0.5 mg/ml at 37°C for 30 min) and proteinase K (final concentration of 0.5 mg/ml at 55°C for 1 h) were treated. Samples were de-crosslinked at 65°C overnight, and ChIP DNA was purified using a ChIP DNA clean and concentrator kit (Zymo).

Muscle ChIP was performed as described previously (Gao et al., 2010). Tissue was chopped into small pieces on the ice and fix solution (50 mM HEPES pH 7.5, 1 mM EDTA pH 8.0, 0.5 mM EGTA, 100 mM NaCl) was added to cross-link the tissue sample. After incubation for 15 min at room temperature, glycine was added as 0.125 M final concentration to stop the reaction. The sample was washed using cold PBS three times followed by homogenizing it in cell lysis buffer (10 mM Tris-HCl pH 8.0, 10 mM NaCl, 0.2% NP40). Cell lysate was centrifuged at 12,000 rpm for 5 min and suspended in nuclear lysis buffer (1% SDS, 10 mM EDTA pH 8.0, 50 mM Tris-HCl pH 8.0). Sonication was performed using Covaris E220 Ultrasonicator (duty cycle:5%, intensity:4, cycle/burst:200, time:120 sec). The resulting chromatin was diluted by 10 fold using dilution buffer (1% Triton X-100, 150 mM NaCl, 2 mM EDTA, 20 mM Tris-HCl pH 8.0). To reduce non-specific binging to beads, the diluted chromatin was mixed with Dynabeads protein A/G for 1 h at 4 °C and the beads were removed prior to incubating chromatin with 2 ug of the appropriate antibodies with Dynabeads protein A/G. After 4 h incubation at 4 °C, beads were washed three times with wash buffer and once with final wash buffer, LiCl Buffer and TE buffer each. Wash buffer contains 1% Triton X-100, 150 mM NaCl, 2 mM EDTA, 20 mM Tris-HCl pH 8.0, 0.1% SDS and final wash buffer contains 500 mM NaCl instead of 150 mM NaCl. The composition of LiCl buffer is 0.25 M LiCl, 1% NP40, 1% deoxycholic acid, 1 mM EDTA, 10 mM Tris-HCl. ChIP DNA was eluted by incubating at 65°C overnight in elution buffer containing 0.25% SDS, 1 mM EDTA, 10 mM Tris-HCl pH7.5. After treatment of proteinase K and RNase A, DNA was purified using ethanol precipitation and MinElute kit (QIAGEN).

Purified DNA (1-5 ng) was used for ChIP-seq library construction with NEBNext ChIP-Seq Library Prep Master Mix Set. ChIP DNA was end-repaired and added with dA tail using Klenow fragment. Sequencing adaptors were ligated to the dA-tailed libraries, and the libraries raging around 270 bp were selected using AMPure XP beads (Beckman Coulter). Size-selected libraries were enriched by PCR with index primers. The quantity and quality of libraries were respectively monitored by library quantification kit (Kapa Biosystems) and Bioanalyzer (Agilent) for 75bp, paired-end Illumina NextSeq.

### RNA-sequencing

RNA was prepared using EZNA total RNA kit (Omega Bio-tek). Total RNA (5 µg) was subjected to ribosomal RNA depletion and ribo-depleted RNA was quantified. Ribo-depleted RNA (20 ng was fragmented and the first strand of cDNA was synthesized using SuperScript III Reverse Transcriptase. Then the second strand was synthesized, and the cDNA was end-repaired and tailed with dA. The end repaired, dA tailed cDNA libraries were ligated with index sequencing adaptors and amplified by PCR. The quality and quantity of the RNA-seq libraries were monitored by Bioanalyzer (Agilent) and library quantification kit (Kapa Biosystems) for 75 bp, paired-end Illumina NextSeq.

### ATAC-sequencing

ATAC-seq was performed as described in (Buenrostro et al., 2013). Nuclei were isolated from 50,000 cells in cold lysis buffer (10 mM Tris-HCl pH 7.4, 10 mM NaCl, 3 mM MgCl_2_, 0.1% NP40, protease inhibitor cocktail). The isolated nuclei were mixed with Nextera (Illumina) TD buffer and Tn5 transposase and incubated at 37°C for 30 min for transposition reaction. The transposed DNA was purified using MinElute PCR Purification Kit (Qiagen) and amplified by PCR with Nextera index primers. The initial 5 cycles of PCR were performed, and an aliquot of the initial reaction was subjected to qPCR to determine additional cycle number of the next PCR. ATAC-seq libraries were purified using MinElute PCR Purification Kit (Qiagen). The quantity and quality of libraries were respectively monitored by library quantification kit (Kapa Biosystems) and Bioanalyzer (Agilent) for 75 bp, paired-end Illumina NextSeq.

### Hi-C

Cells were fixed in PBS containing 1% formaldehyde at RT for 15 min, then quenched by adding glycine at final concentration 0.125 M on ice for 10 min. Dovetail Hi-C libraries were prepared in a similar manner as described previously (Lieberman-Aiden et al., 2009). Briefly, for each library, chromatin was fixed in place with formaldehyde in the nucleus and then extracted. Fixed chromatin was digested with DpnII, the 5’ overhangs filled in with biotinylated nucleotides, and then free blunt ends were ligated. After ligation, crosslinks were reversed and the DNA purified from protein. Purified DNA was treated to remove biotin that was not internal to ligated fragments. The DNA was then sheared to ∼350 bp mean fragment size and sequencing libraries were generated using NEBNext Ultra enzymes and Illumina-compatible adapters. Biotin-containing fragments were isolated using streptavidin beads before PCR enrichment of each library.

### Immunocytochemistry

Cells were washed with sterile PBS and fixed with 4% paraformaldehyde. Fixed cells were permeabilized with 0.5% Triton X-100 in PBS, then blocked with 2% PBA (PBS containing 2% bovine serum albumin) overnight. Primary antibodies were incubated in 2% PBA at RT for 1hr and cells were washed with PBS 3 times. The secondary antibodies (Alexa Fluor 488 Goat Anti-Mouse IgG or Alexa Fluor 568 Goat Anti-Rabbit IgG) were incubated in 2% PBA at RT for 30 min. After PBS washes, nuclei were stained with antifade mounting medium containing DAPI (Vector Laboratories). Immunofluorescence was examined using Olympus Fluoview FV1000 confocal microscope.

### Senescence-associated β-galactosidase assay

Senescence-associated β-galactosidase assay were performed using Senescence β-galactosidase staining kit (Cell Signaling Technology). When cells were not senescent at 4d post-treatment, the standard medium was switched to low serum (0.1% FBS) medium to preserve senescent cells that appeared later. Cells were fixed in 4% paraformaldehyde at RT for 10 min. Cells were washed with PBS and stained in β-galactosidase staining solution containing X-gal (pH 6) at 37°C for overnight in dry incubator. Stained cells were monitored under bright field microscopy.

### Small molecule-driven neuronal reprogramming

Neuronal reprogramming was performed as described in (Li et al., 2015). MEFs were transferred to matrigel-coated plates. When MEFs were confluent, MEF growth medium was switched to Neurobasal Medum containing 1% N2 and 2% B27 supplements, 1% GlutaMAX (Life technologies), 1% penicillin/streptomycin, 100 ng/ml bFGF (STEM CELL), 20 µM ISX9, 100 µM Forskolin, 0.5 µM I-BET151 (CAYMAN CHEM), 20 µM CHIR99021 (LC Laboratories), 2 µM Fasudil and 1 µM SB203580 (Selleckchem). After 2 days, cells were maintained without Fasudil and SB203580. qPCR to detect neuronal gene activation was performed at day 2 after switching to Neurobasal medium, and Tuj1 immunocytochemistry was performed at day 13.

### Adipogenic reprogramming

MEFs or adult fibroblasts were cultured at confluence for 1 day, then the growth medium was switched to the adipogenic differentiation medium (DMEM containing 20% FBS, 1% penicillin/streptomycin, 0.1 mM β-mercaptoethanol, 5 µM dexamethasone, 0.2 mM 3-Isobutyl-1-methylxanthine and 10 µg/ml insulin). Cells were maintained in the differentiation medium for 3 days then subsequently switched to the growth medium containing 10 µg/ml insulin for 4 days. AdipoRed staining was performed at day 7 when lipid droplets were evident in the cells.

### 5-Ethynyl-2’-deoxyuridine (EdU) staining

% EdU-positive cells was measured using the Click-iT® EdU Flow Cytometry Assay Kits (Invitrogen). Briefly,10 μM EdU was added to the culture medium and incubated for 1h. Cells were trypsinized, washed, fixed and permeabilized for Click-iT reaction. EdU+ cells were analyzed using the BD LSR II flow cytometer.

### Western blotting

Cells were rinsed with cold phosphate-buffered saline (PBS) and lysed in NETN (25 mM Tris-HCl (pH 8.0), 1 mM EDTA, 0.15 M NaCl, 0.5% NP-40) containing protease inhibitors (Roche) and phosphatase inhibitors (SIGMA). The cell lysate was briefly sonicated to solubilized insoluble fraction of the cells then clarified by centrifugation. Equal amounts of protein (20–50 μg) were separated on 14-20% SDS-PAGE gel (Biorad) and transferred onto 0.2 μm nitrocellulose membranes. The membranes were blocked with 5% skim milk and incubated with primary antibodies diluted in the range of 1∶10,000-1∶1,000. After wash with TBST (5 mM Tris HCl (pH 7.4), 150 mM NaCl, 0.01% Tween 20) followed by incubation with primary antibodies, bands were visualized by chemiluminescence. For comparison of total histone levels, the same number of cells were lysed in the same volume of lysis buffer.

### Whole-genome sequencing

Genomic DNA was isolated from snap frozen cells or tissues using DNeasy Blood & Tissue Kit. The genomic DNA was fragmented by an ultrasonicator Covaris at 500 bp peak and TruSeq DNA Library Preparation Kit added DNA adaptors to double strand DNA by following the manufacturer’s instructions of Illumina. Deep whole genome sequencing on an Illumina HiSeq X10 platform were performed at BGI.

### RRBS and epigenetic (DNA methylation) clock

RRBS libraries were prepared in two batches. DNA in the first batch was isolated using Quick-DNA Universal kit (Zymo) and in the second batch using E.Z.N.A. Tissue DNA Kit (Omega Bio-tek). 100 μl of 10 mM Tris-HCl buffer was used to elute the samples. Incubation with 2 μl of RNaseA (Life Technologies) was performed for each sample and followed by a purification using Genomic DNA Clean & Concentrator-10 (Zymo). DNA was eluted in 25 μl of TE buffer (10 mM Tris-HCl, 0.1 mM EDTA, pH 8.0). 100ng of each sample, estimated using using a Qubit 2.0 (Life Technologies), was used to prepare RRBS libraries following the previously reported protocol (Petkovich et al., 2017). Libraries included 6-10 samples. The first batch of samples was sequenced on the Illumina HiSeq 2500 platform using 75 bp paired-end sequencing with more than 14 million reads per sample. The second batch was sequenced on the Illumina HiSeq X Ten using 150 paired-end sequencing with more than 32M reads per sample. To compensate for the low complexity of RRBS libraries 10-20% of phiX was spiked in.

Raw reads were filtered and mapped as previously described (Meer et al., 2018). More than 3.6 M CpG sites were covered in each sample and 2.7M were covered in all samples. Data were normalized using ComBat from the SVA package in R. Only CpG sites covered in all samples were considered for DNA methylation clocks application. This resulted in 89 out of 90 sites being covered for the blood DNA methylation clock. Increase of the threshold for the CpG sites coverage decreased the number of the clock sites included in the analysis.

### Histone mass spectrometry

Histone extraction and qMS were performed as previously described (Luense et al., 2016). Acid-extracted histones were propionylated, trypsin-digested and stage-tip desalted with C18 mini-disks. Desalted histone peptides were separated by reversed-phase HPLC on a Thermo Scientific™ EASY-nLC 1000 system. Histone peptide quantified as described (Luense et al., 2016).

### ChIP-sequencing analysis methods

#### Aligning reads

The techniques described for processing the ChIP-seq and ATAC-seq reads are based on ENCODE/Roadmap guidelines with a few modifications (Gjoneska et al., 2015; Landt et al., 2012; Roadmap Epigenomics et al., 2015). The reads were aligned to the mm10 (GRCm38) genome (Cunningham et al., 2019) using Bowtie 2 (Langmead and Salzberg, 2012). The genome fasta files were first indexed and then aligned using the command: bowtie2 -x /directory/with/reference/genome/rootfilename --fast -U /directoryTree/fastq/SAMPLE.fastq -S /directoryTree/fastq/SAMPLE.sam, where SAMPLE was replaced with a unique sample identifier. Following alignment to the genome, the reads were converted from SAM to BAM format (Li et al., 2009). Low quality reads and reads (q < 20) that did not map to the genome were removed. For visualization and peak calling, the bamToBed command line tool was used to convert the BAM files to a modified BED format, called TAGALIGN, which preserved only the read coordinates (Landt et al., 2012; Quinlan and Hall, 2010).

### Spike-in controls

Equal amounts of D. Melanogaster DNA were spiked in to ChIP-seq samples. In addition to aligning to the mouse genome, we aligned reads to the D. Melanogaster dm6 genome (Cunningham et al., 2019). To provide a sense of total ChIP-seq signal strength, the proportion of reads aligning to the dm6 genome were compared to the proportion aligning to the mouse genome (Orlando et al., 2014). To compare Cre and ICE mice, a student’s t-test was used on those proportions.

### Visualizing read coverage

For visualizing individual samples, the genomecov tool within BedTools was used to convert from the BED format to a BEDGRAPH format (Quinlan and Hall, 2010). Finally, the BEDGRAPH file was converted to the more efficient BIGWIG format using the UCSC command line tool bedGraphToBigWig (Kent et al., 2010; Speir et al., 2016). The information was uploaded to the NCBI sequence read archive and SRA files representing the raw reads, the BAM file representing the aligned reads, and the BigWig files of read coverage across the genome.

### Visualizing signal relative to background

Macs2 bdgcmp command was used to calculate the signal to noise ratio for every position in the genome for each combination of histone modification and experimental condition (Cre and ICE) (Feng et al., 2012). The BEDGRAPH file was converted to the more efficient BIGWIG format using the UCSC command line tool bedGraphToBigWig (Speir et al., 2016).

### Quantification and statistical analysis

#### Peak Calling

For each epigenetic measurement (H3K27ac, H3K56ac, H3K27me3, ATAC-seq) and input samples, the TAGALIGN files were merged using the unix command zcat and sorted using the unix sort command according to the chromosome using the start position. For each histone modification, MACS2 was used to call the peaks relative to the input control: macs2 callpeak -t H3K27ac.tagAlign.gz -f BED -c input.tagAlign.gz -n H3K27ac_signal -g mm -p 1e-2 --nomodel -- extsize 73 -B –SPMR (Feng et al., 2012). For ATAC-Seq, no input control was used. We removed peaks with significance (signal relative to noise) of p > 10^−5^. Peaks that fell into the ENCODE blacklist regions were removed (Landt et al., 2012). The output of the program is a BED file with peak coordinates for the mm10 version of the mouse genome.

### Peak annotation

Peaks are annotated based on their mapping to the nearest transcription start site, which was performed using BEDTools closestBed command (Quinlan and Hall, 2010) based on ENSEMBL gene annotations, GRCm38 version 79 (Cunningham et al., 2019).

### Counting reads across peaks

For each histone modification and ATAC-seq sample, the reads from each experiment are counted in the called peaks using featureCounts in the subread package (Liao et al., 2013). To perform the counting the peak BED file were converted to SAF format.

### Differential peaks between Cre and ICE

The negative binomial model in the DESeq2 R package was used to identify differential peaks between the CRE and ICE mice (Love et al., 2014). For MEF experiments, we used a stringent threshold of adjusted p < 0.01. For muscle ChIP-Seq experiments, very few peaks attained significance levels at that cutoff. Therefore, we restricted our analysis to looking at the group of peaks enriched at p < 0.01. The varianceStabilizingTransformation function in the DESeq2 package was used to normalize the read counts per peak. Sex chromosomes were excluded from analyses due to the inconsistency of the sexes of the MEFs.

### Transcription factor binding sites

First, BED files of the ICE-higher, ICE-lower, and the total set of peaks were converted in the FASTA formats of the corresponding genome sequence using the twobit2fa program available from the UCSC genome browser command line tools (Speir et al., 2016). The transcription factor binding site analysis was conducted using the Meme suite (Bailey, 2011; Khan et al., 2018). Within the MEME suite, the FIMO program was used to identify individual transcription factor binding sites within peaks (MA0139.1 for CTCF). The ame program within the meme suite was used to identify enriched transcription factor binding site motifs in the high and lower regulatory elements of each histone modification relative to the background of all peaks for that histone modification. For each enrichment calculation, we used the Wilcox rank sum test method and fisher’s exact test. To be stringent, we required that both statistical tests return a significant value (adjusted p<0.05) for the site to be reported. In the initial set of results, large groups of very similar transcription factor binding site motifs were shown to be enriched. To remove that redundancy, we clustered the JASPAR transcription factor binding site motifs using the STAMP tool (Mahony et al., 2007; Mahony and Benos, 2007). Transcription factor binding site motifs with a distance of < 10^−5^ were grouped together. We report the most significance score for each group of transcription factor binding site motifs.

### Super-enhancer (SE) comparison

We compared the differential peaks to mouse super-enhancers in dbSUPER (Khan and Zhang, 2016). To facilitate that comparison, we mapped super-enhancers to the mm10 version of the genome using liftover (Speir et al., 2016). For the two binary variables, differential histone modification and presence/absence of a super-enhancer signal in that cell type, a hypergeometric test was used to calculate the significance of the overlap. We also report the fold enrichment of the observed overlap relative the number of overlapping peaks expected if super-enhancer presence was independent of differential histone modification.

### ChIP-sequencing / RNA-sequencing comparisons

To compare the ChIP-seq and RNA-Seq differences between the CRE and the ICE mice, a matrix of RNA-Seq read counts was obtained (see other section). To minimize noise in the comparison, DESeq2 was used to recalculate the log2 fold difference in expression between the Cre and ICE mice. For each histone modification and the ATAC-seq experiment, we counted the number of reads per sample at the promoter (+/− 1 kb from transcription start). For each histone modification and each set of differential peaks (ICE higher and ICE lower), the distribution log_2_ fold change in gene expression between ICE and Cre is examined. The significance of the relationship of each of these peak categories to gene expression is determined using a student’s t-test.

### Basline ChIP-sequencing / RNA-sequencing comparisons

To test whether the differences in gene expression were function of the histone modifications present at baseline or the differences in histone modification between Cre and ICE, we used the quantification of histone modifications at the promoter regions. DESeq2 (Love et al., 2014) was used to model the read count:

read count ∼ Intercept + ICE_cells. The model was able to estimate baseline levels of the histone modification (Intercept) along with the log2 fold changes associated with each histone modification relative to the WCE control. In parallel, a negative binomial model was used to identify the genes that were differentially expressed in the ICE MEFs relative to the CRE (p adj. < 0.01; log2FC >= 1; 391 higher ICE; 818 lower in ICE). We used a t-test to identify whether the significantly ICE higher or ICE lower genes showed a bias in histone modification at the promoter, both at baseline (intercept term) and in response to DNA damage in the ICE model.

### ChIP-seq metaplots and heatmaps

Metaplots and heatmaps were produced using deepTools version 3.0.1 (Ramirez et al., 2016). Intermediate matrix files were generated by applying computeMatrix (scale-regions mode) to BIGWIG files over genomic loci in BED format. plotProfile and plotHeatmap functions were applied to the matrix files to generate output data used to graph each metaplot and heatmap.

### Gene ontology of ChIP-seq differential peaks

Gene ontology analysis for ChIP-seq were performed using Genomic Regions Enrichment of Annotations Tool (GREAT) (McLean et al., 2010). Genomic coordinates of differential ChIP-seq regions and all ChIP-seq peaks were used as test regions and background regions, respectively. GO biological processes were ranked by HyperFdrQ and only GO terms made up of at least 5 genes were included. ChIP-seq data were also analyzed using ChIP-Enrich (Welch et al., 2014). GO terms with at least 5 genes were ranked by FDR.

### RNA-seq analysis

Reads were aligned with hisat2 v2.1.0 (Kim et al., 2015) to the Ensembl GRCm38 primary assembly using splice junctions from the Ensembl release 84 annotation. Paired read counts were quantified using featureCounts v1.6.4 (Liao et al., 2014) using reads with a MAPQ >=20. Diffentially-expressed genes for each pairwise comparison were identified with edgeR v3.26 (Robinson et al., 2010), testing only genes with at least 0.1 counts-per-million (CPM) in at least three samples. Gene ontology analysis of differentially-expressed genes was performed with MetaCore from Clarivate Analytics.

### Hi-C analysis

Paired-end reads were aligned with bwa mem (v0.7.17) (Li and Durbin, 2009) using the options -S -P. Interaction were parsed and deduplicated with pairsamtools (v0.0.1). Pairwise interaction frequencies were binned in 50-kb nonoverlapping windows and intra-chromosome interaction frequencies were normalized by dividing each interaction by the average number of interactions observed with that distance. Normalized interaction matrices were binned with smoothMat (Yang et al., 2017). Matrix pearson’s correlations were calculated in R v 3.6.1 (Bunn, 2008, 2010) and used to perform a principal component analysis. The sign of the first eigenvector for each chromosome was adjusted to correlate with GC content, and were smoothed with loess smoothing using 1 megabase windows.

### Whole-genome sequencing analysis

Whole-Genome raw sequencing reads from paired-end library was quality-controlled with FastQC and subsequently mapped to the reference genome GRCm38/mm10 (mm10) using the Burrows–Wheeler Alignment (BWA-MEM, version 0.7.17) (Li et al., 2009). A paired-end mapping strategy with default parameters was utilized. After mapping, the reads were sorted and converted into binary alignment format (BAM) via Sequence Alignment/Map tools (SAMtools, version 1.9). The best practices recommended by the Broad Institute for variant calling were then followed (Van der Auwera et al., 2013). The sorted binary alignments underwent post-processing to remove duplicates via Picard’s MarkDuplicates (v.2.01; http://broadinstitute.github.io/picard) before germline variants were identified using Genome Analysis Toolkit (GATK; v. 3.7) HaplotypeCaller (McKenna et al., 2010).

### Podocyte density p57 and PAS representative images

Podocyte density was quantitated following staining for p57 on formalin fixed, paraffin-embedded, 4 µm kidney sections as previously described (Ohse et al., 2010; Schneider et al., 2017; Zhang et al., 2013). Briefly, Histoclear (National Diagnostics, Atlanta, GA) was used to deparaffinize kidney sections, followed by rehydration using graded 100%, 95%, and 70% ethanol baths. Next, antigen retrieval was performed using 10 mM EDTA pH 6.0. Endogenous peroxidase activity was blocked with 3% v/v H2O2. Non-specific antibody binding was blocked using a 5% non-fat milk in PBS. Rabbit polyclonal p57 antibody (Santa Cruz) was diluted 1:800 in 1% BSA in PBS, applied to the sections, and incubated overnight at 4°C. Rabbit-on-rodent HRP polymer (Biocare Medical, Concord, CA) was applied and incubated at room temperature for 45 min. Diaminobenzidine (DAB) (Sigma-Aldrich, St. Louis, MO) with 0.05% NiCl (Sigma-Aldrich) was used to detect staining. Slides used for podocyte density were not counterstained in order to improve quantitation sensitivity. For representative images, counterstaining was performed with periodic acid–Schiff. Sections were placed in 0.5% periodic acid (Sigma-Aldrich), washed in ddH2O, incubated for 10 min with Schiff’s Reagent (Sigma-Aldrich), washed in 0.5% sodium metabisulfate (Sigma-Aldrich) and incubated with hematoxylin (Sigma-Aldrich). Tissue was dehydrated in 95% and 100% ethanol baths, followed by Histoclear and Histomount (National Diagnostics). Podocyte density was quantitated according to the correction factor method from single histological sections, as previous reported (Venkatareddy et al., 2014). An average of 119 (±14.1) glomeruli for ICE mice and 139 (±16.1) glomeruli for Cre mice were quantified.

### Glomerular Injury

Organized matrix accumulation was detected on paraffin-embedded tissue by Jones’ basement membrane stain (Silver Stain) performed by the University of Washington Pathology Research Services Laboratory following standard protocols (Luna, 1968). Silver stained slides were quantitated according to the criteria presented in figure 6G. An average of 156 (±10.8) glomeruli for ICE mice and 187 (±6.36) glomeruli for Cre mice were quantified.

### Parietal epithelial cell to mesenchymal transition

Parietal epithelial cells (PECs) were stained for alpha-smooth muscle actin (α-SMA) in order to determine epithelial-mesenchymal transition (EMT) as described above. Non-specific antibody binding was blocked using Background buster (Accurate Chemical & Scientific Corporation, Westbury, NY). Rabbit polyclonal α-SMA antibody (Abcam) was diluted 1:400 in 1% BSA in PBS, applied to the sections, and incubated overnight at 4°C. Detection was performed as described above. Quantification was performed by counting the number of glomeruli with α-SMA staining in PECs as previously described (Schneider et al., 2017). An average of 110 (±6.22) glomeruli for ICE mice and 122 (±19.3) glomeruli for Cre mice were quantified.

### Microscopy and imaging for kidney

Imaging and quantification were performed on a Leica DMI400B microscope and an EVOS FL Cell Imaging System. ImageJ 1.51 (NIH) was used to measure podocyte density.

## Supplemental Information

Table S1. Primers used in this study, Related to Figures 1, 3, 4 and 5

Table S2. QC results of MEF ChIP-seq, Related to Figures 2, S2 and S3

Table S3. Differential MEF ChIP-seq peaks, Related to Figures 2, 3 and 4

Table S4. GO analysis of differential MEF ChIP-seq peaks, Related to Figures 3 and 4

Table S5. TF binding motifs in differential MEF ChIP-seq peaks, Related to Figure S4

Table S6. Differential genes and GO analysis of MEF RNA-seq, Related to Figures 3 and 5

Table S7. Differential genes and OG analysis of adult fibroblast RNA-seq, Related to Figure 5

Table S8. QC results of muscle ChIP-seq, Related to Figures 6

Table S9. Differential muscle ChIP-seq peaks, Related to Figure 6

---

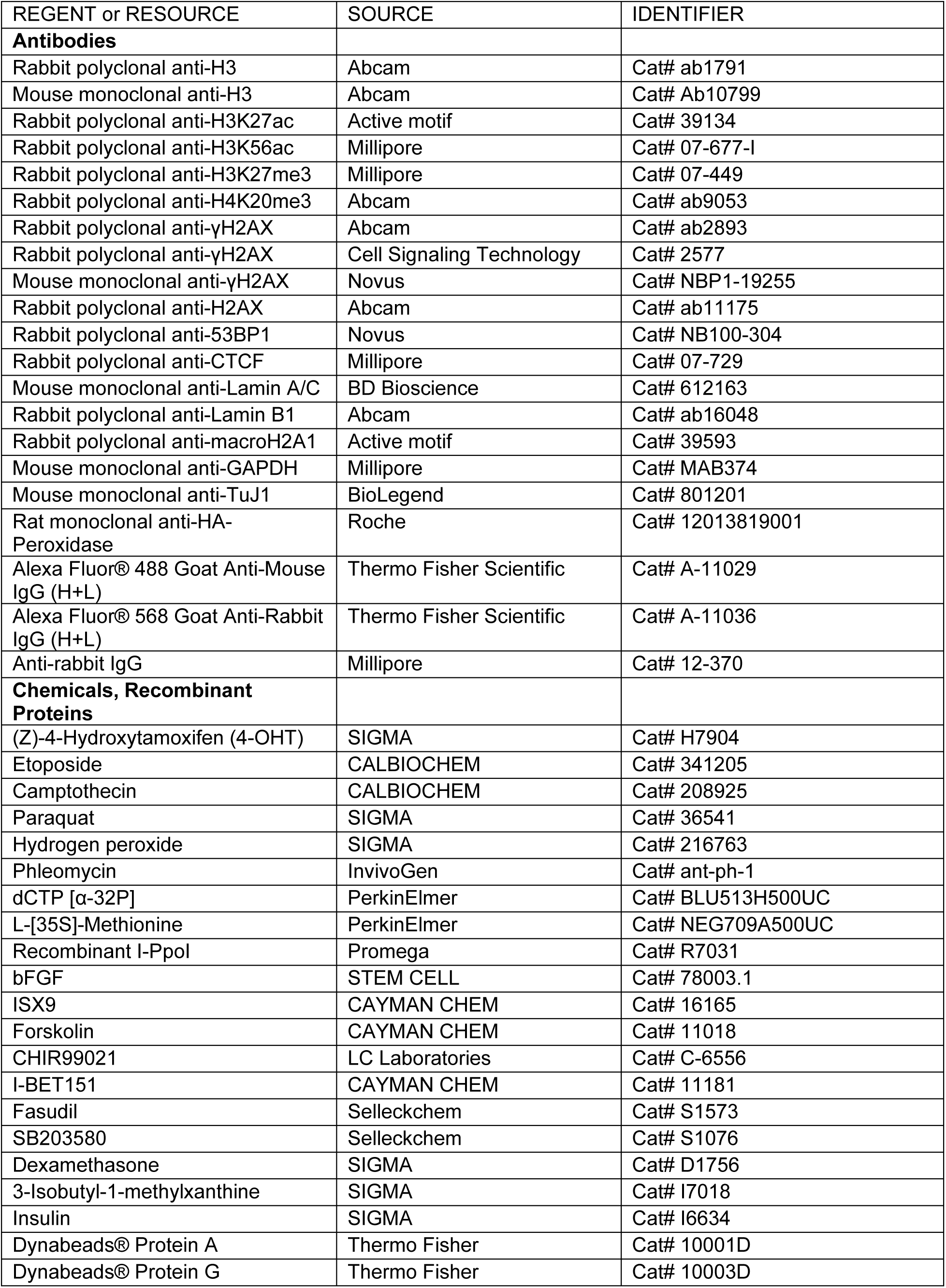
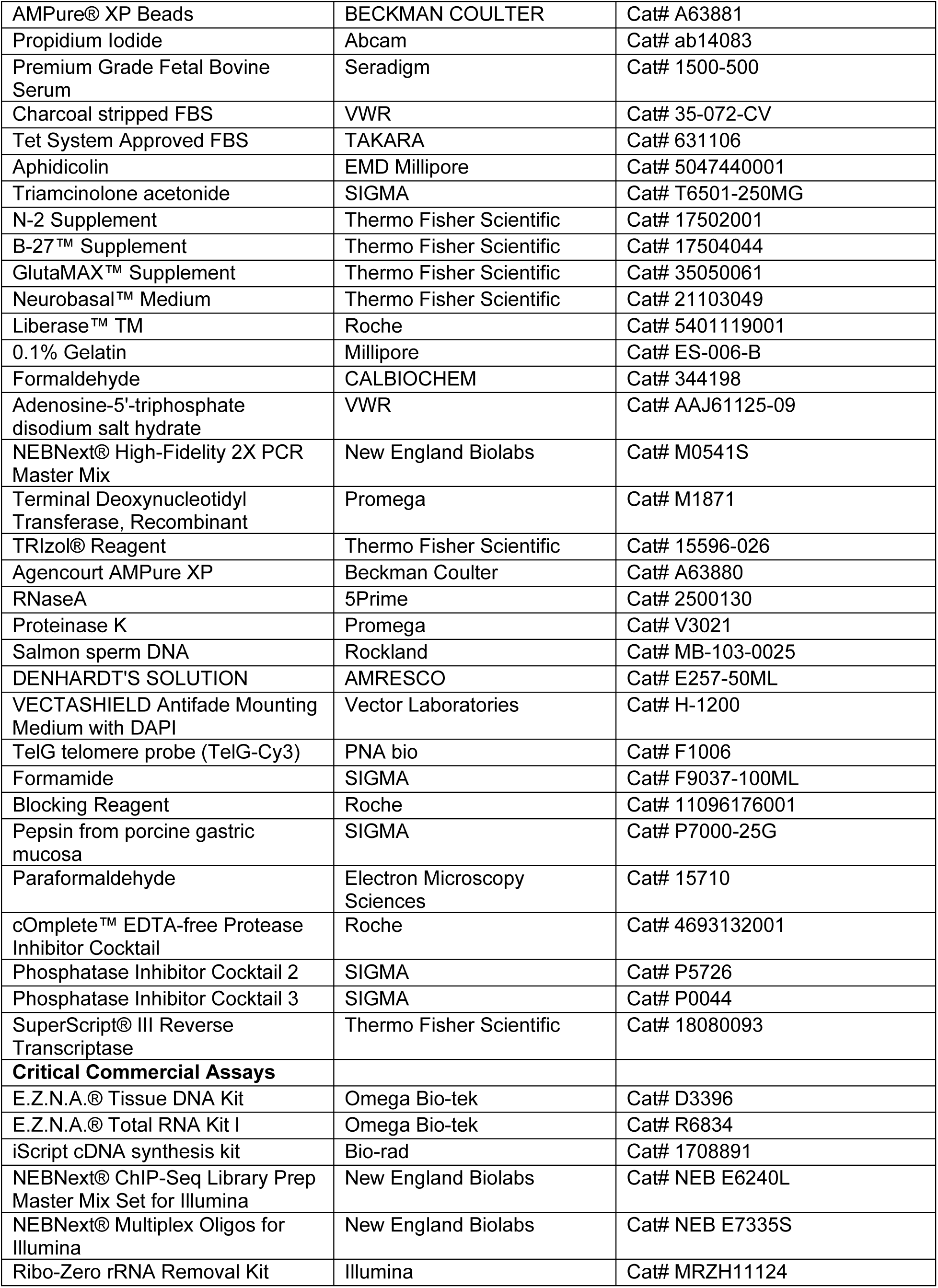
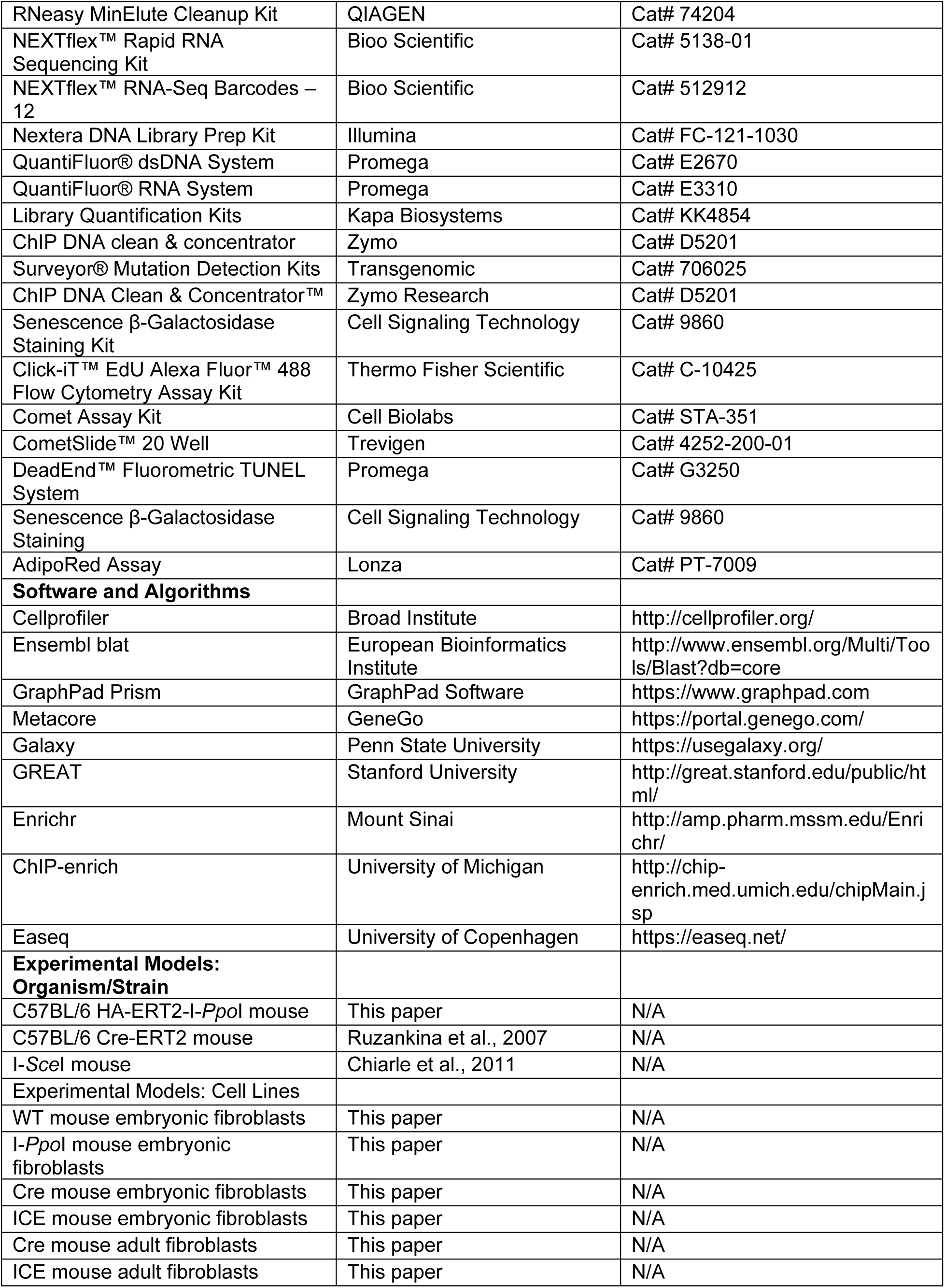
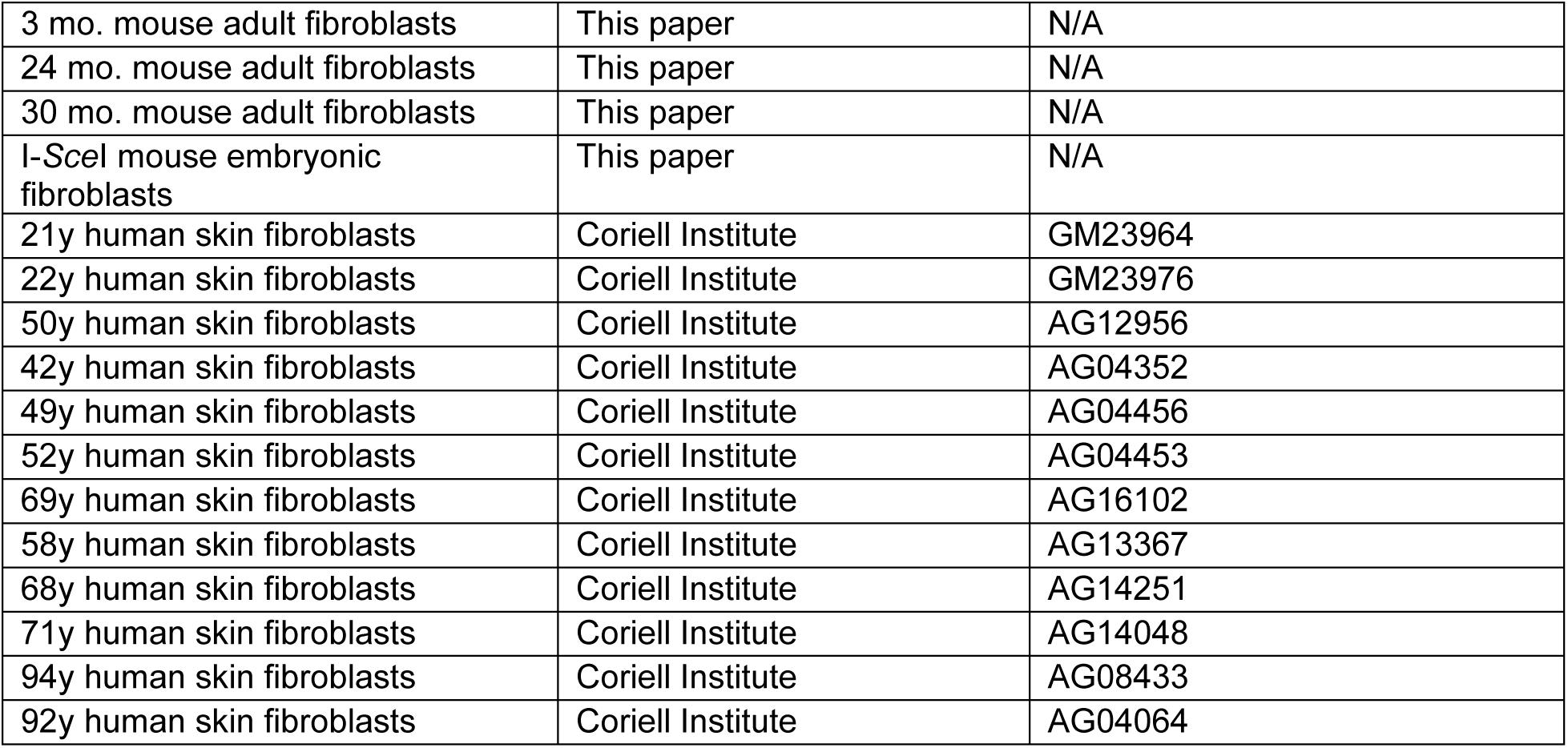

| REAGENT or RESOURCE | SOURCE | IDENTIFIER |
| --- | --- | --- |
| **Antibodies** |  |  |
| Rabbit polyclonal anti-H3 | Abcam | Cat# ab1791 |
| Mouse monoclonal anti-H3 | Abcam | Cat# Ab10799 |
| Rabbit polyclonal anti-H3K27ac | Active motif | Cat# 39134 |
| Rabbit polyclonal anti-H3K56ac | Millipore | Cat# 07-677-I |
| Rabbit polyclonal anti-H3K27me3 | Millipore | Cat# 07-449 |
| Rabbit polyclonal anti-H4K20me3 | Abcam | Cat# ab9053 |
| Rabbit polyclonal anti-γH2AX | Abcam | Cat# ab2893 |
| Rabbit polyclonal anti-γH2AX | Cell Signaling Technology | Cat# 2577 |
| Mouse monoclonal anti-γH2AX | Novus | Cat# NBP1-19255 |
| Rabbit polyclonal anti-H2AX | Abcam | Cat# ab11175 |
| Rabbit polyclonal anti-53BP1 | Novus | Cat# NB100-304 |
| Rabbit polyclonal anti-CTCF | Millipore | Cat# 07-729 |
| Mouse monoclonal anti-Lamin A/C | BD Bioscience | Cat# 612163 |
| Rabbit polyclonal anti-Lamin B1 | Abcam | Cat# ab16048 |
| Rabbit polyclonal anti-macroH2A1 | Active motif | Cat# 39593 |
| Mouse monoclonal anti-GAPDH | Millipore | Cat# MAB374 |
| Mouse monoclonal anti-TuJ1 | BioLegend | Cat# 801201 |
| Rat monoclonal anti-HA-Peroxidase | Roche | Cat# 12013819001 |
| Alexa Fluor® 488 Goat Anti-Mouse IgG (H+L) | Thermo Fisher Scientific | Cat# A-11029 |
| Alexa Fluor® 568 Goat Anti-Rabbit IgG (H+L) | Thermo Fisher Scientific | Cat# A-11036 |
| Anti-rabbit IgG | Millipore | Cat# 12-370 |
| **Chemicals, Recombinant Proteins** |  |  |
| (Z)-4-Hydroxytamoxifen (4-OHT) | SIGMA | Cat# H7904 |
| Etoposide | CALBIOCHEM | Cat# 341205 |
| Camptothecin | CALBIOCHEM | Cat# 208925 |
| Paraquat | SIGMA | Cat# 36541 |
| Hydrogen peroxide | SIGMA | Cat# 216763 |
| Phleomycin | InvivoGen | Cat# ant-ph-1 |
| dCTP [α-32P] | PerkinElmer | Cat# BLU513H500UC |
| L-[35S]-Methionine | PerkinElmer | Cat# NEG709A500UC |
| Recombinant I-Ppol | Promega | Cat# R7031 |
| bFGF | STEM CELL | Cat# 78003.1 |
| ISX9 | CAYMAN CHEM | Cat# 16165 |
| Forskolin | CAYMAN CHEM | Cat# 11018 |
| CHIR99021 | LC Laboratories | Cat# C-6556 |
| I-BET151 | CAYMAN CHEM | Cat# 11181 |
| Fasudil | Selleckchem | Cat# S1573 |
| SB203580 | Selleckchem | Cat# S1076 |
| Dexamethasone | SIGMA | Cat# D1756 |
| 3-Isobutyl-1-methylxanthine | SIGMA | Cat# I7018 |
| Insulin | SIGMA | Cat# I6634 |
| Dynabeads® Protein A | Thermo Fisher | Cat# 10001D |
| Dynabeads® Protein G | Thermo Fisher | Cat# 10003D |
| AMPure® XP Beads | BECKMAN COULTER | Cat# A63881 |
| Propidium Iodide | Abcam | Cat# ab14083 |
| Premium Grade Fetal Bovine Serum | Seradigm | Cat# 1500-500 |
| Charcoal stripped FBS | VWR | Cat# 35-072-CV |
| Tet System Approved FBS | TAKARA | Cat# 631106 |
| Aphidicolin | EMD Millipore | Cat# 5047440001 |
| Triamcinolone acetonide | SIGMA | Cat# T6501-250MG |
| N-2 Supplement | Thermo Fisher Scientific | Cat# 17502001 |
| B-27™ Supplement | Thermo Fisher Scientific | Cat# 17504044 |
| GlutaMAX™ Supplement | Thermo Fisher Scientific | Cat# 35050061 |
| Neurobasal™ Medium | Thermo Fisher Scientific | Cat# 21103049 |
| Liberase™ TM | Roche | Cat# 5401119001 |
| 0.1% Gelatin | Millipore | Cat# ES-006-B |
| Formaldehyde | CALBIOCHEM | Cat# 344198 |
| Adenosine-5'-triphosphate disodium salt hydrate | VWR | Cat# AAJ61125-09 |
| NEBNext® High-Fidelity 2X PCR Master Mix | New England Biolabs | Cat# M0541S |
| Terminal Deoxynucleotidyl Transferase, Recombinant | Promega | Cat# M1871 |
| TRIzol® Reagent | Thermo Fisher Scientific | Cat# 15596-026 |
| Agencourt AMPure XP | Beckman Coulter | Cat# A63880 |
| RNaseA | 5Prime | Cat# 2500130 |
| Proteinase K | Promega | Cat# V3021 |
| Salmon sperm DNA | Rockland | Cat# MB-103-0025 |
| DENHARDT'S SOLUTION | AMRESCO | Cat# E257-50ML |
| VECTASHIELD Antifade Mounting Medium with DAPI | Vector Laboratories | Cat# H-1200 |
| TelG telomere probe (TelG-Cy3) | PNA bio | Cat# F1006 |
| Formamide | SIGMA | Cat# F9037-100ML |
| Blocking Reagent | Roche | Cat# 11096176001 |
| Pepsin from porcine gastric mucosa | SIGMA | Cat# P7000-25G |
| Paraformaldehyde | Electron Microscopy Sciences | Cat# 15710 |
| cOmplete™ EDTA-free Protease Inhibitor Cocktail | Roche | Cat# 4693132001 |
| Phosphatase Inhibitor Cocktail 2 | SIGMA | Cat# P5726 |
| Phosphatase Inhibitor Cocktail 3 | SIGMA | Cat# P0044 |
| SuperScript® III Reverse Transcriptase | Thermo Fisher Scientific | Cat# 18080093 |
| **Critical Commercial Assays** |  |  |
| E.Z.N.A.® Tissue DNA Kit | Omega Bio-tek | Cat# D3396 |
| E.Z.N.A.® Total RNA Kit I | Omega Bio-tek | Cat# R6834 |
| iScript cDNA synthesis kit | Bio-rad | Cat# 1708891 |
| NEBNext® ChIP-Seq Library Prep Master Mix Set for Illumina | New England Biolabs | Cat# NEB E6240L |
| NEBNext® Multiplex Oligos for Illumina | New England Biolabs | Cat# NEB E7335S |
| Ribo-Zero rRNA Removal Kit | Illumina | Cat# MRZH11124 |
| RNeasy MinElute Cleanup Kit | QIAGEN | Cat# 74204 |
| NEXTflex™ Rapid RNA Sequencing Kit | Bioo Scientific | Cat# 5138-01 |
| NEXTflex™ RNA-Seq Barcodes – 12 | Bioo Scientific | Cat# 512912 |
| Nextera DNA Library Prep Kit | Illumina | Cat# FC-121-1030 |
| QuantiFluor® dsDNA System | Promega | Cat# E2670 |
| QuantiFluor® RNA System | Promega | Cat# E3310 |
| Library Quantification Kits | Kapa Biosystems | Cat# KK4854 |
| ChIP DNA clean & concentrator | Zymo | Cat# D5201 |
| Surveyor® Mutation Detection Kits | Transgenomic | Cat# 706025 |
| ChIP DNA Clean & Concentrator™ | Zymo Research | Cat# D5201 |
| Senescence β-Galactosidase Staining Kit | Cell Signaling Technology | Cat# 9860 |
| Click-iT™ EdU Alexa Fluor™ 488 Flow Cytometry Assay Kit | Thermo Fisher Scientific | Cat# C-10425 |
| Comet Assay Kit | Cell Biolabs | Cat# STA-351 |
| CometSlide™ 20 Well | Trevigen | Cat# 4252-200-01 |
| DeadEnd™ Fluorometric TUNEL System | Promega | Cat# G3250 |
| Senescence β-Galactosidase Staining | Cell Signaling Technology | Cat# 9860 |
| AdipoRed Assay | Lonza | Cat# PT-7009 |
| **Software and Algorithms** |  |  |
| Cellprofiler | Broad Institute | http://cellprofiler.org/ |
| Ensembl blat | European Bioinformatics Institute | http://www.ensembl.org/Multi/Tools/Blast?db=core |
| GraphPad Prism | GraphPad Software | https://www.graphpad.com |
| Metacore | GeneGo | https://portal.genego.com/ |
| Galaxy | Penn State University | https://usegalaxy.org/ |
| GREAT | Stanford University | http://great.stanford.edu/public/html/ |
| Enrichr | Mount Sinai | http://amp.pharm.mssm.edu/Enrichr/ |
| ChIP-enrich | University of Michigan | http://chip-enrich.med.umich.edu/chipMain.jsp |
| Easeq | University of Copenhagen | https://easeq.net/ |
| **Experimental Models: Organism/Strain** |  |  |
| C57BL/6 HA-ERT2-I-*Ppol* mouse | This paper | N/A |
| C57BL/6 Cre-ERT2 mouse | Ruzankina et al., 2007 | N/A |
| I-SceI mouse | Chiarle et al., 2011 | N/A |
| **Experimental Models: Cell Lines** |  |  |
| WT mouse embryonic fibroblasts | This paper | N/A |
| I-*Ppol* mouse embryonic fibroblasts | This paper | N/A |
| Cre mouse embryonic fibroblasts | This paper | N/A |
| ICE mouse embryonic fibroblasts | This paper | N/A |
| Cre mouse adult fibroblasts | This paper | N/A |
| ICE mouse adult fibroblasts | This paper | N/A |
| 3 mo. mouse adult fibroblasts | This paper | N/A |
| 24 mo. mouse adult fibroblasts | This paper | N/A |
| 30 mo. mouse adult fibroblasts | This paper | N/A |
| I-Scel mouse embryonic fibroblasts | This paper | N/A |
| 21y human skin fibroblasts | Coriell Institute | GM23964 |
| 22y human skin fibroblasts | Coriell Institute | GM23976 |
| 50y human skin fibroblasts | Coriell Institute | AG12956 |
| 42y human skin fibroblasts | Coriell Institute | AG04352 |
| 49y human skin fibroblasts | Coriell Institute | AG04456 |
| 52y human skin fibroblasts | Coriell Institute | AG04453 |
| 69y human skin fibroblasts | Coriell Institute | AG16102 |
| 58y human skin fibroblasts | Coriell Institute | AG13367 |
| 68y human skin fibroblasts | Coriell Institute | AG14251 |
| 71y human skin fibroblasts | Coriell Institute | AG14048 |
| 94y human skin fibroblasts | Coriell Institute | AG08433 |
| 92y human skin fibroblasts | Coriell Institute | AG04064 |

**Figure S1.**
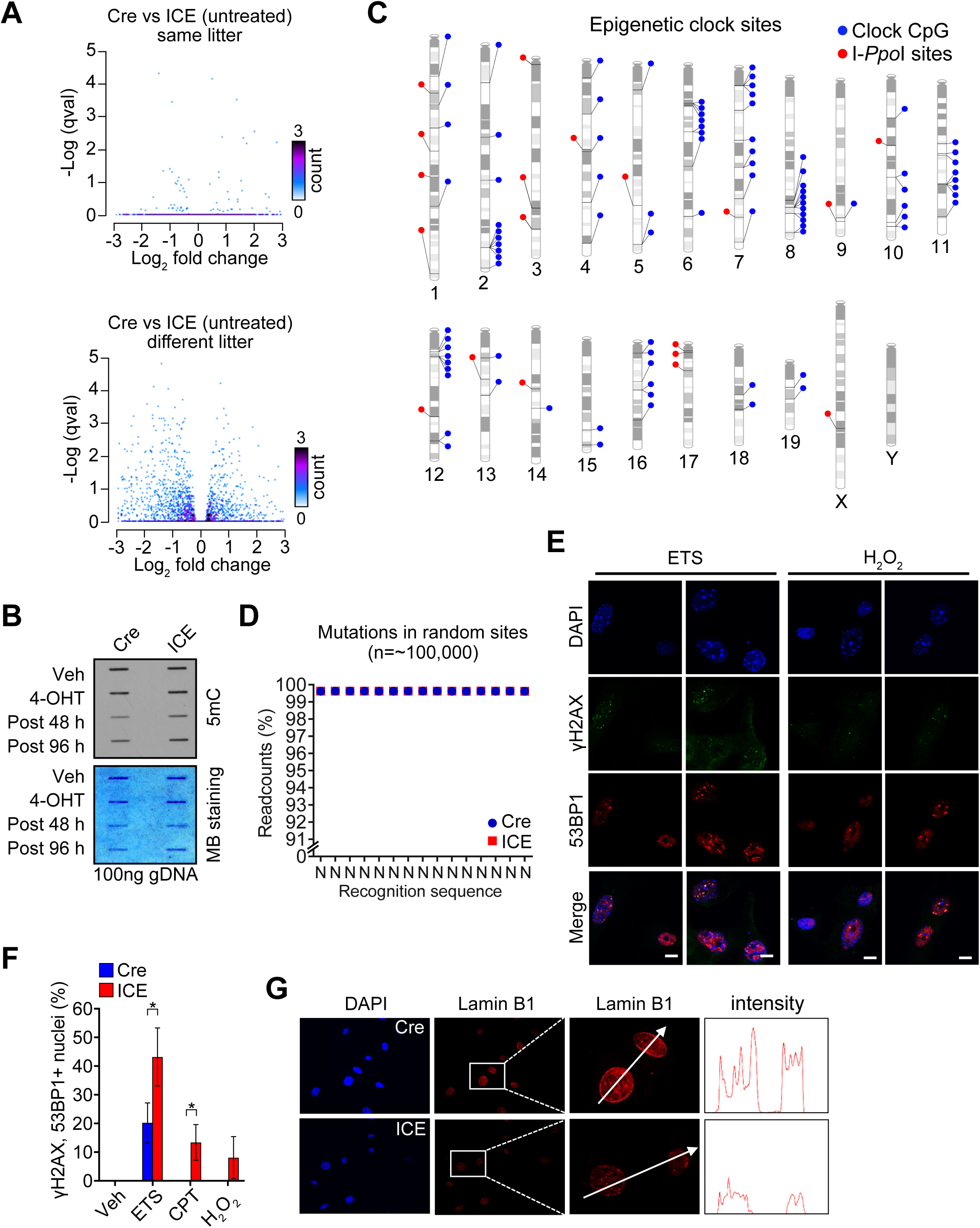
A Cell-based System to Study the Effect of DSBs on the Epigenome, Related to Figure 1. (A) RNA-seq volcano plots (Cre vs. ICE) of cells from the same or different litters before 4-OHT treatment. (B) Slot blots to compare global 5mC levels. Methylene blue (MB) staining showed total genomic DNA used in each sample. (C) Genomic distribution of I-*Ppo*I canonical sites and all 90 CpG sites of the mouse epigenetic clock. (D) Percent non-mutated sequences of ∼100,000 random sites in post-treated ICE cells assessed by deep sequencing (>50x). (E and F) Immunostaining of DNA damage markers γH2AX and 53BP1 in post-treated ICE cells with and without exposure to the DNA damaging agents (ETS, etoposide; CPT, camptothecin; H_2_O_2_, hydrogen peroxide). Two-tailed Student’s *t* test. Scale bar, 10 µm. (G) Immunostaining of Lamin B1 in post-treated ICE cells. Data are mean (n=3) ± SD. *p < 0.05; ***p< 0.001.

**Figure S2.**
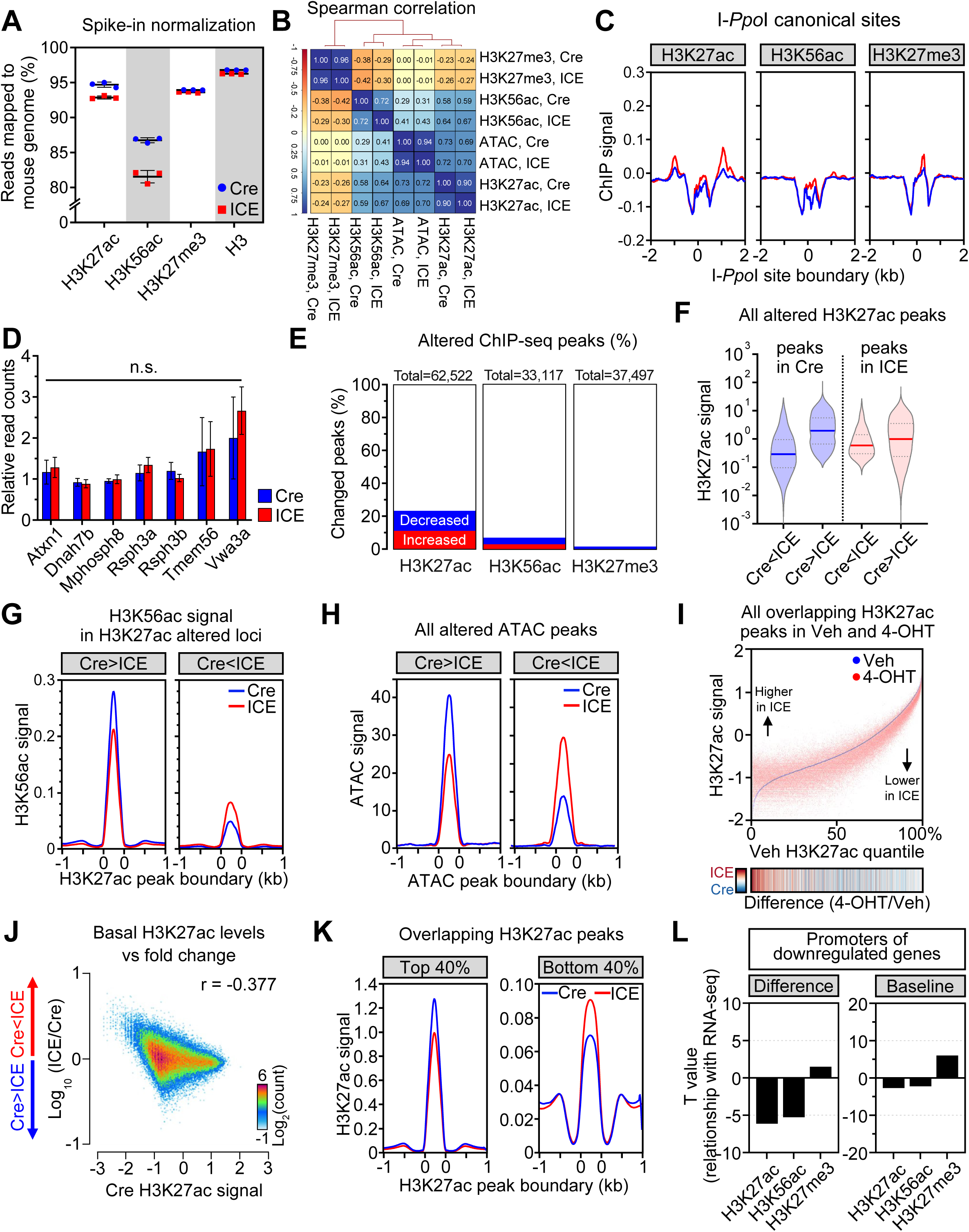
Altered Epigenetic Landscapes in ICE Cells, Related to Figure 2. (A) Spike-in normalization of ChIP-seq data. (B) Spearman correlation matrix between ChIP-seq and ATAC-seq experiments. (C) Aggregation plots of I-*Ppo*I canonical sites. (D) Normalized RNA-seq read counts of genes overlapped with I-*Ppo*I canonical sites. Two-tailed Student’s *t* test. Data are mean (n=3) ± SD. n.s.: p > 0.05 (E) Percent changed peaks of each histone modification. (F) Violin plots of changed H3K27ac peak signals. (G) Aggregation plots of H3K56ac signal in H3K27ac changed regions. (H) Aggregation plots of ATAC signal in ATAC changed regions. (I) Genome-wide changes of H3K27ac in ICE cells compared to Veh-treated ICE controls. Heatmap of 4-OHT/Veh. (J) Correlation between H3K27ac signals in Cre and fold change between Cre and ICE. (K) Aggregation plots of H3K27ac signal in top 40% or bottom 40% quantile. (L) Correlation between histone modifications and mRNA levels. Difference, Cre vs. ICE; Baseline, signal in Cre.

**Figure S3.**
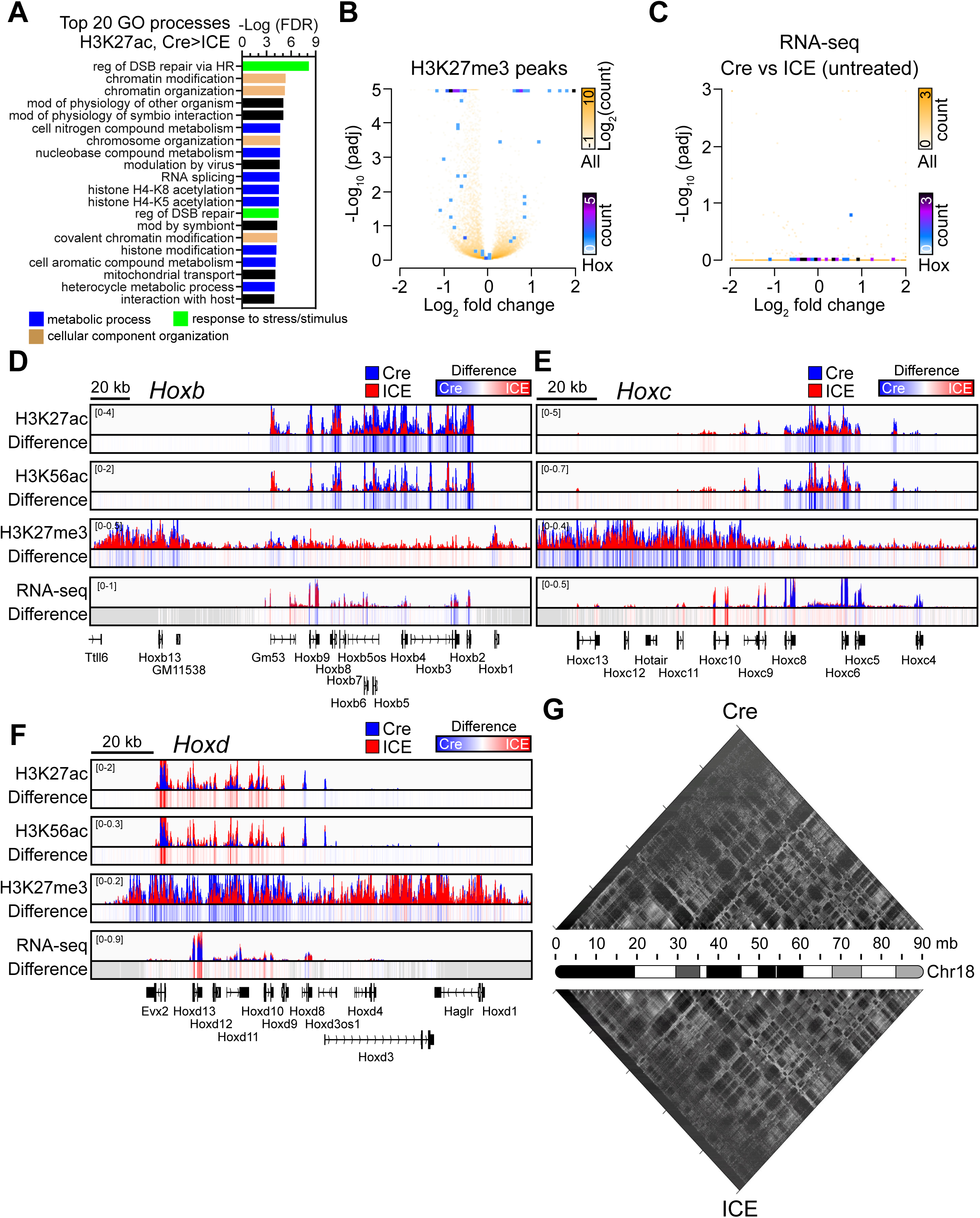
Dysregulation of *Hox* Loci in Epigenetically Aged ICE Cells, Related to Figure 3. (A) Gene Ontology analysis of histone ChIP-seq data ordered by top 20 processes enriched in H3K27ac-decreased regions (padj < 0.01). (B and C) Volcano plot of H3K27me3 peaks and RNA-seq. All genes and *Hox* genes shown white to yellow and blue to purple, respectively. (D - F) ChIP-seq track of histone modifications and mRNA levels across the *Hox* loci of post-treated ICE cells. (G) Normalized Hi-C contact matrices of chromosome 18 at 50 kb resolution.

**Figure S4.**
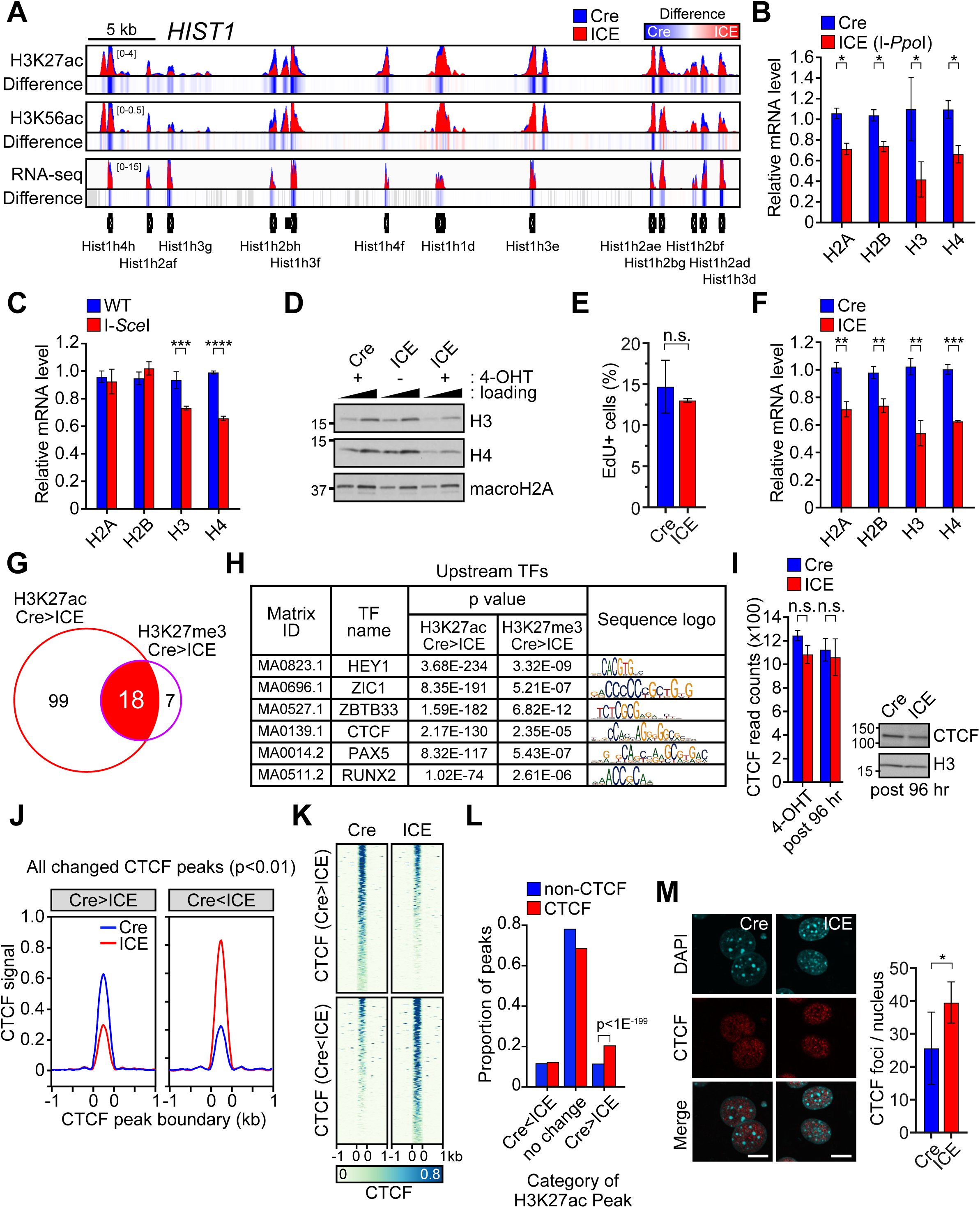
Dysregulation of *Hist1* Loci and CTCF Binding in Epigenetically Aged ICE Cells, Related to Figure 3. (A) ChIP-seq track of histone modifications and mRNA levels across the *Hist1* locus of post-treated ICE cells. (B and C) qPCR analysis of canonical histone genes, H2A, H2B, H3 and H4, in post-treated cells cut with either I-*Ppo*I or I-*Sce*I homing endonucleases. Two-tailed Student’s *t* test. (D) Wester blotting of histone H3 and H4 in post-treated ICE cells. macroH2A serves as a loading control. (E) Percent 5-Ethynyl-2’-deoxyuridine (EdU)-positive cells. (F) qPCR analysis of H2A, H2B, H3 and H4, in post-treated ICE cells upon cell cycle arrest by aphidicolin treatment. Two-tailed Student’s *t* test. (G) Number of transcription factor binding sites enriched with H3K27ac or H3K27me3 ChIP-seq data. (H) List of transcription factors involved in development and DNA repair. (I) Normalized RNA-seq read counts and protein levels of CTCF in 4-OHT-treated or post-treated ICE cells. Two-tailed Student’s *t* test. (J and K) Aggregation plots and heatmaps of CTCF signal in CTCF changed regions (p < 0.01). (L) Proportion of CTCF peaks found in H3K27ac changed regions in post-treated ICE cells. (M) Immunostaining and quantification of CTCF foci in post-treated ICE cells. Scale bar, 10 µm. Two-tailed Student’s *t* test. Data are mean (n≥3) ± SD. n.s.: p > 0.05; *p < 0.05; **p < 0.01; **** p < 0.0001.

**Figure S5.**
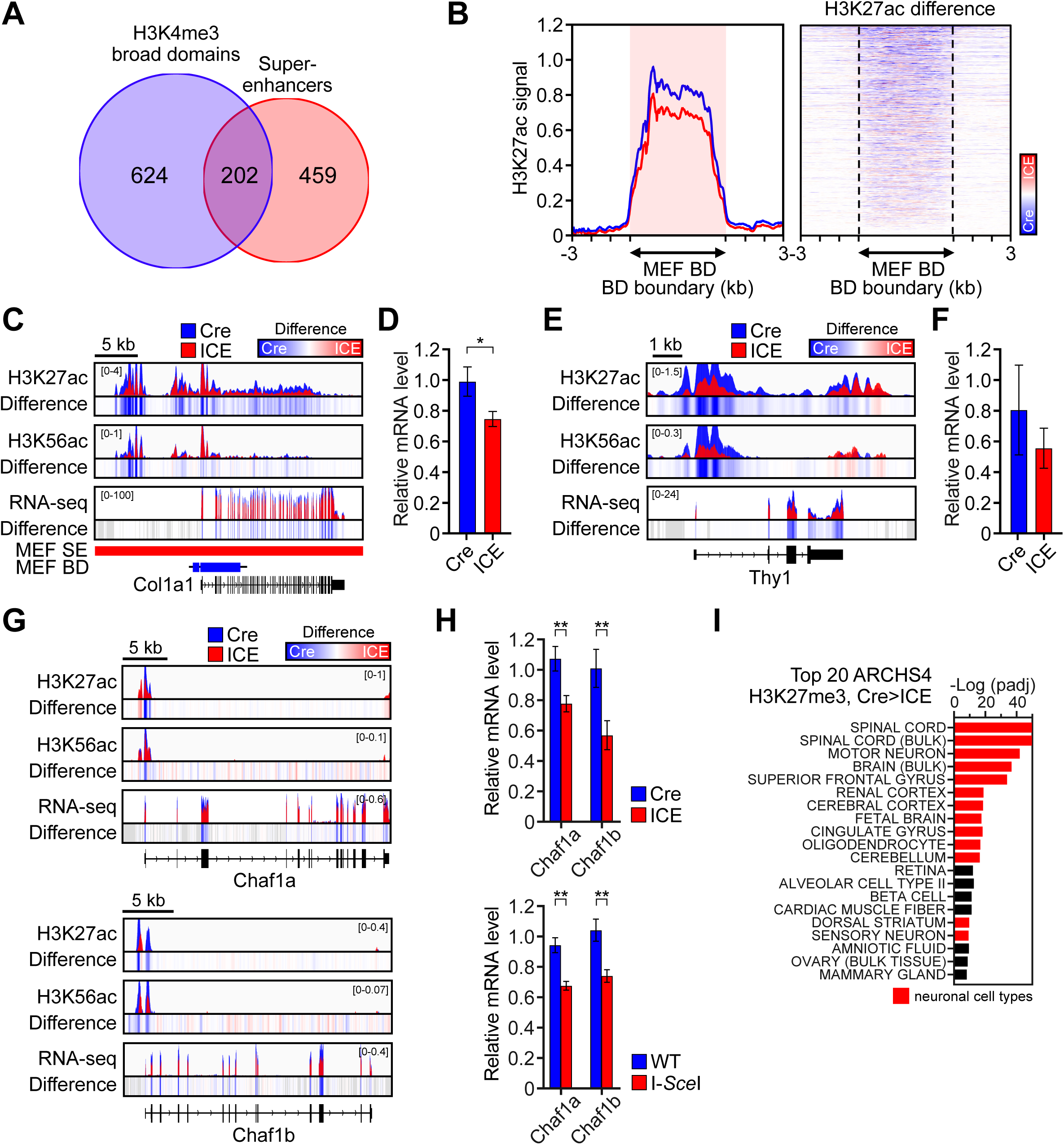
DSBs Induce Epigenetic Changes at Genes That Control Cellular Identity, Related to Figure 4. (A) Number of the top 5% broadest H3K4me3 domains and super-enhancers in MEFs. (B) Aggregation plots and heatmaps of H3K27ac signals in top 5% MEF broadest H3K4me3 domains. (C - F) ChIP-seq track of histone modifications and mRNA levels across *Col1a1* and *Thy1*, in post-treated ICE cells. (G) ChIP-seq track of histone modifications and mRNA levels across the *Chaf1a* and *Chaf1b* in post-treated ICE cells. (H) qPCR analysis of *Chaf1a* and *Chaf1b* in post-treated cells cut with either I-*Ppo*I or I-*Sce*I homing endonucleases. Two-tailed Student’s *t* test. (I) Human tissue types of transcriptional profiles that overlap decreased H3K27me3 regions (padj < 0.05) in epigenetically aged ICE cells. Red represents neuronal cell types. Data are mean (n≥3) ± SD. *p < 0.05; **p < 0.01.

**Figure S6.**
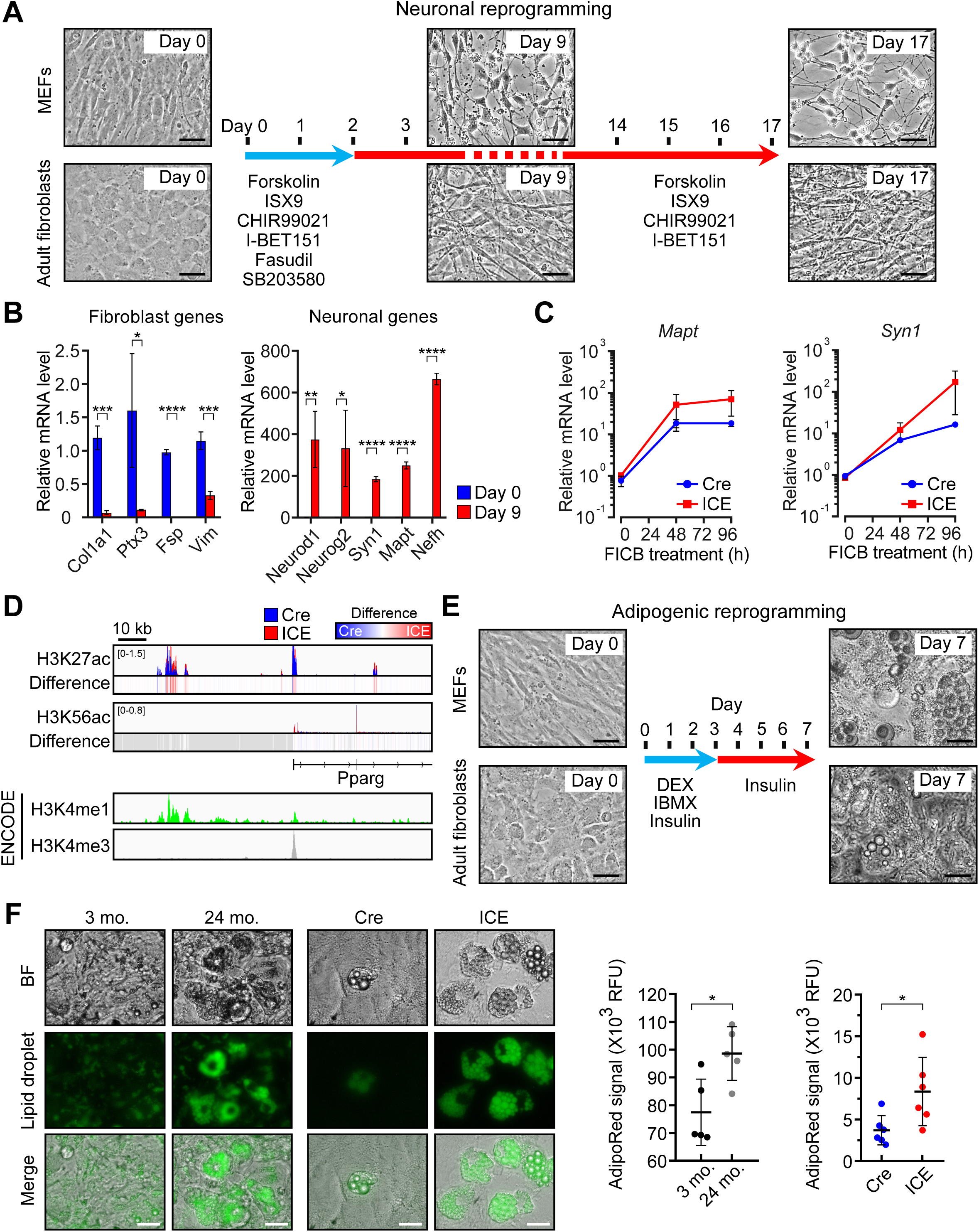
Increased Plasticity of Cellular Identity in Epigenetically Aged ICE Cells, Related to Figure 5. (A) Representative images of wild type MEFs or adult fibroblasts from 3 month-old mice during small molecule-derived neuronal reprogramming. Scale bar, 50 µm. (B) qPCR analysis of fibroblast genes and neuronal genes at day 0 and day 9. Two-tailed Student’s *t* test. (C) Time-course analysis of mRNA levels of *Mapt* and *Syn1* during neuronal reprogramming. (D) ChIP-seq track of histone modifications upstream regulatory regions of *Pparg* of post-treated ICE cells. The promoter and the enhancer of *Pparg* were respectively enriched with H3K4me3 or H3K4me1 in ENCODE ChIP-seq data. (E) Representative images of wild type MEFs or adult fibroblasts from 3 month-old mice during adipogenic reprogramming. Scale bar, 50 µm. (F) Representative images and quantification of lipid droplets in post-treated ICE MEFs or adult fibroblasts from wild type 3 or 24 month-old mice. Scale bar, 50 µm. Two-tailed Student’s *t* test. Data are mean (n≥3) ± SD. *p < 0.05; **p < 0.01; ***p < 0.001; ****p < 0.0001.

**Figure S7.**
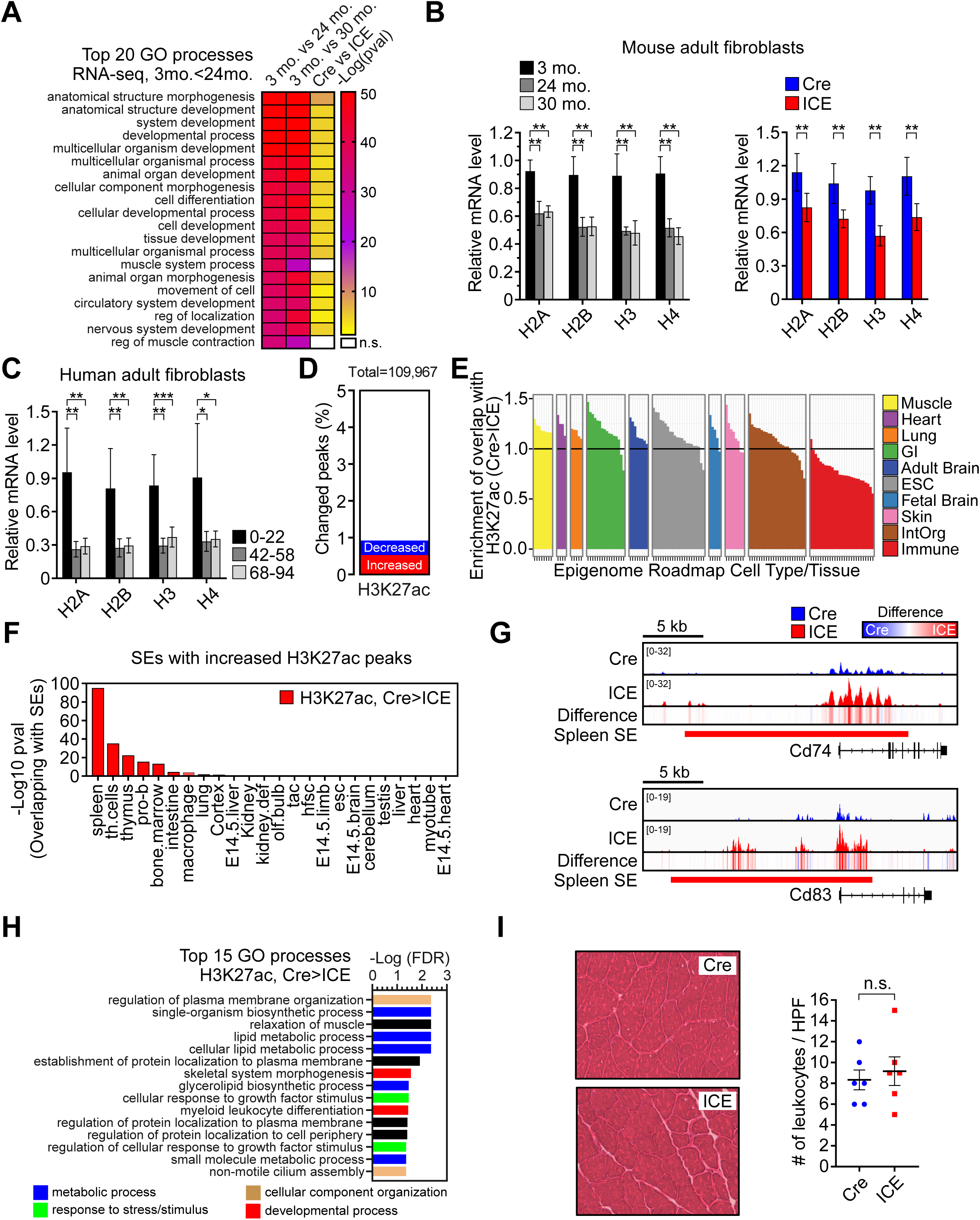
Cells and Tissues of ICE Mice Mimic Those from Old Wildtype Mice, Related to Figure 6. (A) Gene Ontology analysis of RNA-seq data ordered by top 20 processes enriched in up-regulated genes in fibroblasts from 24 month-old mice (padj < 0.05, FC > 1.5). (B and C) mRNA levels of histone genes in adult fibroblasts from wild type 3, 24, and 30 month-old mice, 6 month-old ICE mice and human adult fibroblasts of different ages. One-way ANOVA-Bonferroni (B, left and C), two-tailed Student’s *t* test (B, right). (D) Percent changed peaks of H3K27ac in 10-month post-treated ICE muscle. (E) Comparison of H3K27ac decreased regions (p < 0.01) to epigenome roadmap data from different human tissue types. (F) Super-enhancers (SEs) in different cell types that overlap with regions with increased H3K27ac signals (Cre>ICE in ICE muscle). (G) ChIP-seq track of histone modifications across *Cd74* and *Cd83* of 10-month post-treated ICE muscle. (H) Gene Ontology analysis of histone ChIP-seq data ordered by top 20 processes enriched in H3K27ac decreased regions in 10-month post-treated ICE muscle (p < 0.01). (I) H&E staining and quantification of infiltrated leukocytes in 10-month post-treated ICE muscle. Data are mean (n≥3) ± SD. *p < 0.05; **p < 0.01; ***p < 0.001.

